# Oncogenic *PIK3CA* corrupts growth factor signaling specificity

**DOI:** 10.1101/2023.12.23.573207

**Authors:** R.R. Madsen, A. Le Marois, O. Mruk, M. Voliotis, S. Yin, J. Sufi, X. Qin, S.J. Zhao, J. Gorczynska, D. Morelli, L. Davidson, E. Sahai, V.I. Korolchuk, C.J. Tape, B. Vanhaesebroeck

## Abstract

Pathological activation of the PI3K/AKT pathway is among the most frequent defects in human cancer and is also the cause of rare overgrowth disorders. Yet, there is currently no systematic understanding of the quantitative flow of information within PI3K/AKT signaling and how it is perturbed by disease-causing mutations. Here, we develop scalable, single-cell approaches for systematic analyses of signal processing within the PI3K pathway, enabling precise calculations of its information transfer for different growth factors. Using genetically-engineered human cell models with allele dose-dependent expression of *PIK3CA^H1047R^*, we show that this oncogene is not a simple, constitutive pathway activator but a context-dependent modulator of extracellular signal transfer. *PIK3CA^H1047R^*reduces information transmission downstream of IGF1 while selectively enhancing EGF-induced signaling and transcriptional responses. This leads to a gross reduction in signaling specificity, akin to “blurred” signal perception. The associated increase in signaling heterogeneity promotes phenotypic diversity in a human cervical cancer cell line model and in human induced pluripotent stem cells. Collectively, these findings and the accompanying methodological advances lay the foundations for a systematic mapping of the quantitative mechanisms of PI3K/AKT-dependent signal processing and phenotypic control in health and disease.

**One-sentence summary:** Single-cell signaling and information theoretic analyses reveal that oncogenic PI3K/AKT activation leads to a gross reduction in signaling specificity, context-dependent EGF response amplification as well as increased phenotypic heterogeneity.

## INTRODUCTION

The class IA phosphoinositide 3-kinase (PI3K)/AKT pathway is essential for cellular and organismal homeostasis. It is used for signal transduction downstream of most if not all growth factors (GFs) as well as many hormones and cytokines. The pathway also represents a key therapeutic target due to its frequent hyperactivation across human cancers. This is often due to mutations in *PIK3CA*, the gene encoding the p110α catalytic subunit of the PI3Kα isoform. Based on early cellular studies (1–3), common cancer-associated *PIK3CA* mutations such as *PIK3CA^H1047R^*are often regarded as constitutive “on” switches, or activators, of the pathway. Consequently, therapeutic targeting of aberrant PI3K/AKT activation in this context has focused on pathway switch-off (4). However, the efficacy of this approach is often limited by the toxicity of PI3K/AKT inhibition in healthy cells and tissues treated with high doses of PI3K/AKT inhibitors. This is true not only in cancer but also in the non-cancerous *PIK3CA*-related overgrowth spectrum (PROS) of congenital disorders caused by an identical spectrum of activating *PIK3CA* mutations as in cancer (5,6).

The “switch” view of the impact of activating *PIK3CA* mutations and the resulting therapeutic limitations reflect a more general, critical gap in the current knowledge of PI3K/AKT signaling. Specifically, there is limited understanding of how quantitative, dynamic patterns of PI3K/AKT activation are used by cells to specify (i.e., encode) the identity of the myriad environmental signals sensed by this pathway (7,8). It therefore also remains unknown if and how disease-causing mutations in PI3K/AKT pathway components may perturb this temporal code. For example, corruption of dynamic signal encoding has been documented in the related RAS/MAPK signaling cascade in response to certain oncogenic BRAF mutations and targeted inhibitors (9).

This type of quantitative mapping of the input-output relationships in the PI3K/AKT pathway is technically very challenging. It requires capture of multimodal biochemical responses with high temporal resolution and quantitative precision (7,8). Moreover, unlike conventional protein phosphorylation cascades, the key first step in PI3K pathway activation is the generation of the plasma membrane-localized lipid second messenger phosphatidylinositol-3,4,5-trisphosphate (PIP_3_), and its derivative PI(3,4)P_2_. The detection of these low-abundance lipids presents a technical challenge, and thus the vast majority of studies of oncogenic PI3K signaling do not feature direct evaluation of this critical first signal encoding step in PI3K pathway activation, focusing instead on bulk measurements of downstream effector responses (8).

Lastly, a breakthrough in translating our currently semi-quantitative view of the PI3K/AKT pathway into a fully quantitative framework is contingent upon access to systematic measurements of PI3K/AKT signaling at the single-cell level (7,8). This is important for two reasons. First, the fidelity of information transfer within a system depends both on the strength of the signal and on its uncertainty, or variability (10). The latter refers to the biochemical response heterogeneity that would typically be observed at the level of individual cells in an otherwise homogenous cell population. This concept is at the core of mathematical information theoretic analyses of signaling pathways, a powerful approach pioneered by Levchenko and colleagues to study quantitative signaling fidelity (11,12). Second, robust approaches to measurements of PI3K/AKT signaling heterogeneity at the single-cell level are required for mapping signaling thresholds to phenotypic decision making according to a probabilistic framework (8,13,14). This contrasts with the conventional view of deterministic outputs downstream of PI3K/AKT pathway activation.

Here, we first set out to address the technical limitations that preclude systematic studies of single-cell PI3K/AKT signaling at scale. Following extensive benchmarking of available PIP_3_/PI(3,4)P_2_ biosensors and optimization of available live-cell and mass-cytometry-based protocols, we have developed robust experimental and analytical approaches for studies of single-cell PI3K/AKT biology. We then used these approaches to study quantitative signal transfer in cell models with allele dose-dependent expression of *PIK3CA^H1047R^*. We discovered that this oncogene corrupts the fidelity of signal transmission in cervical cancer (HeLa) cells and in induced pluripotent stem cells. The associated increase in PI3K/AKT and RAS/MAPK signaling heterogeneity manifested phenotypically in the emergence of co-existing cell states. Our findings and methodological advances provide the basis for quantitative mapping of PI3K/AKT-dependent signal processing and phenotypic control in health and disease.

## RESULTS

### Optimized workflow for PI3K activity measurements at the plasma membrane

For quantitative studies of the immediate phosphoinositide lipid outputs of PI3K activation in individual cells and at high temporal resolution, we established a robust, semi-automated live-cell imaging pipeline (**Fig. S1A**). Using total internal reflection fluorescence (TIRF) and the small-molecule PI3Kα activator 1938 (15) to monitor PI3Kα outputs specifically at the plasma membrane, we first systematically benchmarked the quantitative fidelity (dynamic range, technical variability) of several PH domain-based phosphoinositide biosensors (**Fig. S1B**). These included the PH domain of BTK (with or without the adjacent TH domain (16,17)), a tandem-dimer version of the PH domain of ARNO (with modifications to minimize interactions with other proteins (18)), and the PH domain of AKT2 (of note, this is not the full-length protein to avoid internalization independent of PIP_3_ binding (19)). For consistent comparisons, all biosensors were expressed from the same plasmid backbone, featuring a GFP tag at the C-terminus and a nuclear export sequence at the N-terminus (**Fig. S1B**). As control for specificity, all experiments with wild-type biosensors also featured co-expression of an mCherry-tagged version of each construct, with an arginine-to-alanine mutation that ablates phosphoinositide binding (**Fig. S1B**) (20).

The PH domain of AKT2 consistently performed as an optimal biosensor, based on the dynamic range, reproducibility across experiments, low sensitivity to technical noise, and rapid response upon activation as well as inhibition of PI3Kα (**Fig. S1C, S1D**). We note, however, that while this measures the total PIP_3_/PI(3,4)P_2_ output of PI3K activation at the plasma membrane, the original N terminal-tagged version of the PH-TH of BTK would be a better option for studies that seek to selectively study PIP_3_ independent of PI(3,4)P_2_ (**Fig. S1E**).

Our final optimized TIRF-based workflow for PI3K activity measurements allowed profiling of up to 4 different cellular conditions (e.g., genotypes) and 60 single cells per experiment, with live-cell measurements taken every 70 sec over 60 min while exposing the cells to controlled perturbations, giving rise to more than 3000 individual data points.

### Temporal measurements of class IA PI3K activation identify conserved, dynamic encoding of growth factor signals

Using a set of independent cellular model systems (human HeLa cervical cancer cells, human A549 lung adenocarcinoma cells, immortalized mouse embryonic fibroblasts) with or without endogenous functional PI3Kα, we next tested the hypothesis that the identity of different growth factors is captured in the cellular dynamics of PIP_3_/PI(3,4)P_2_. Using saturating doses of IGF1 and EGF in serum-free medium, we observed remarkably consistent responses that suggested conservation of the dynamic signal encoding of these growth factors across different cell models and species (**Fig. 1**). At the population level, both IGF1- and EGF-induced PIP_3_/PI(3,4)P_2_ reporter responses exhibited a characteristic overshoot, with a peak within the first 10 min of stimulation, followed by a sustained quasi steady-state above baseline. The key difference between the two growth factors was in the response amplitude, with PI3Kα wild-type cell lines reaching a peak PIP_3_/PI(3,4)P_2_ fold-change of ∼1.55 for IGF1 and ∼1.25 for EGF **(Fig. 1)**.

**Figure 1.**
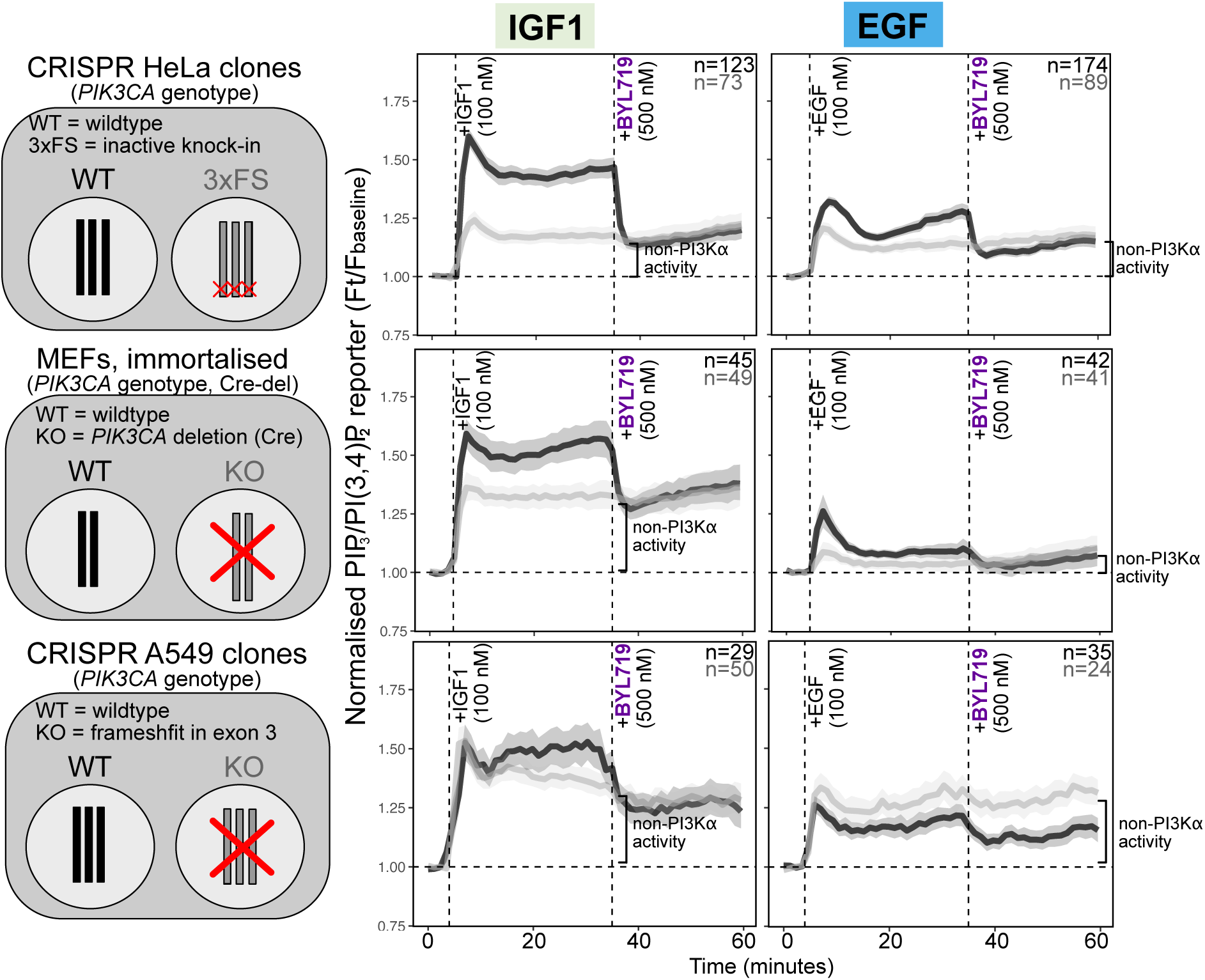
IGF1 and EGF induce stereotypical PIP_3_/PI(3,4)P_2_ signaling dynamics. TIRF microscopy measurements of dynamic IGF1- and EGF-induced PIP_3_/PI(3,4)P_2_ levels in live HeLa, MEF or A549 cells with wild-type or loss-of-function *PIK3CA* as indicated. The cells were serum-starved for 3 h prior to stimulation with either saturating doses (100 nM) of IGF1 or EGF, followed by PI3Kα-selective inhibition with 500 nM BYL719. The traces represent the mean PH_AKT2_ reporter fold-change relative to baseline (the median signal of the first four time points). The shading signifies bootstrapped 95% confidence intervals of the mean. The number (n) of single cells for each genotype is indicated on the plots. For wild-type (WT) HeLa cells, two independent CRISPR/Cas9 clones were used, with and without silent mutations. The 3x FS HeLa cells originate from a single CRISPR/Cas9 clone, engineered with a frameshift mutation in all three *PIK3CA* alleles (see also Fig. S2). The MEFs were from polyclonal cultures established from mice with the respective *PIK3CA* genotypes, followed by immortalization *in vitro* (82). The A549 cells were from a single CRISPR/Cas9 clone per genotype. HeLa datasets for IGF1 and EGF are from 6 and 7 independent experiments, respectively. MEF and A549 IGF1 and EGF data are from 3 independent experiments each. Non-PI3Kα activity refers to the class IA PI3K activity that remains following pharmacological inhibition of PI3Kα.

The combined genetic and pharmacological (BYL719) inactivation of PI3Kα in these experiments also revealed cell type- and growth factor-specific quantitative differences in the contribution of the PI3Kα isoform to each growth factor response (**Fig. 1**). For example, in HeLa cells, approximately 60% and 50% of the IGF1 and EGF response, respectively, was mediated by PI3Kα. In mouse embryonic fibroblasts, PI3Kα contributed 40% of the IGF1 response and up to 60% of the peak EGF response yet only up to 50% of the sustained EGF-induced PIP_3_/PI(3,4)P_2_ reporter response. This would therefore suggest that the observed stereotypical IGF1- and EGF-dependent PIP_3_/PI(3,4)P_2_ response patterns are robust to the relative contribution of individual class IA PI3K isoforms (**Fig. 1**).

Collectively, these data identify conserved dynamic PI3K-dependent encoding of IGF1 and EGF, with high temporal and isoform-specific resolution.

### Oncogenic *PIK3CA^H1047R^* reduces the PIP_3_/PI(3,4)P_2_ information capacity of IGF1

We next tested whether the dynamic signal encoding of growth factor identity is equally robust to the expression of *PIK3CA^H1047R^*, one of the most commonly observed oncogenic PI3Kα mutations in cancer and PROS. Allele dose-dependent, endogenous expression of this variant was engineered in HeLa cells, using CRISPR/Cas9 (**Fig. S2A, S2B**). We chose this cell model due to its low baseline PI3K/AKT signaling (**Fig. S2**), absence of pathway-specific mutations, in-depth characterization at multiple biological levels (transcriptomics, proteomics (21)), experimental tractability and, in particular, its cervical cancer origin. Genomic profiling of human cervical tumors has revealed this cancer to be among the most enriched for multiple *PIK3CA* mutations in *cis* or *trans* (22,23). We therefore reasoned that HeLa cells may allow to capture allele dose-dependent effects of *PIK3CA^H1047R^*on quantitative signal transfer. So far, such allele dose-dependent effects have only been studied mechanistically in a developmental model system (24,25).

Several quality control assays were applied to all final CRISPR/Cas9 clones to identify possible confounders. Assays included whole-exome sequencing (**Fig. S2C**), transcriptomics (**Fig. S2D**) and candidate-based mRNA and protein expression evaluations (**Fig. S2E, S2F, S2G**). None of these assays revealed any systematic differences across the different clones except for the desired knock-in of *PIK3CA^H1047R^* and evidence for an associated yet subtle baseline PI3K/AKT pathway activation by immunoblotting. This is important because it enables to study the consequences of the oncogenic perturbation on signaling response independent of widespread transcriptional changes that could modify the topology of the relevant signaling networks.

To assess dynamic signal encoding of IGF1 and EGF as a function of *PIK3CA* genotype, cells were stimulated with one of three different doses (1 nM, 10 nM, 100 nM) of each growth factor, and PIP_3_/PI(3,4)P_2_ responses were captured using TIRF microscopy as described above (**Fig. 2A**). The resulting temporal measurements of PI3K activity at the single-cell level were processed for mathematical information-theoretic analyses of trajectory responses (26), resulting in formal quantification of the fidelity of dose-dependent signal transfer through PI3K activation. Given three different stimulus doses for each growth factor, the theoretical maximum information capacity captured in the PIP_3_/PI(3,4)P_2_ response would be 1.58 bits (log2(3)), corresponding to the case where the PIP_3_/PI(3,4)P_2_ response alone is sufficient to distinguish between the different doses of the growth factor perfectly. While this is unlikely to be reached given technical noise, values that are substantially lower than 1.58 would imply that the PIP_3_/PI(3,4)P_2_ response alone is not sufficient to distinguish the different doses of each growth factor with high certainty.

**Figure 2.**
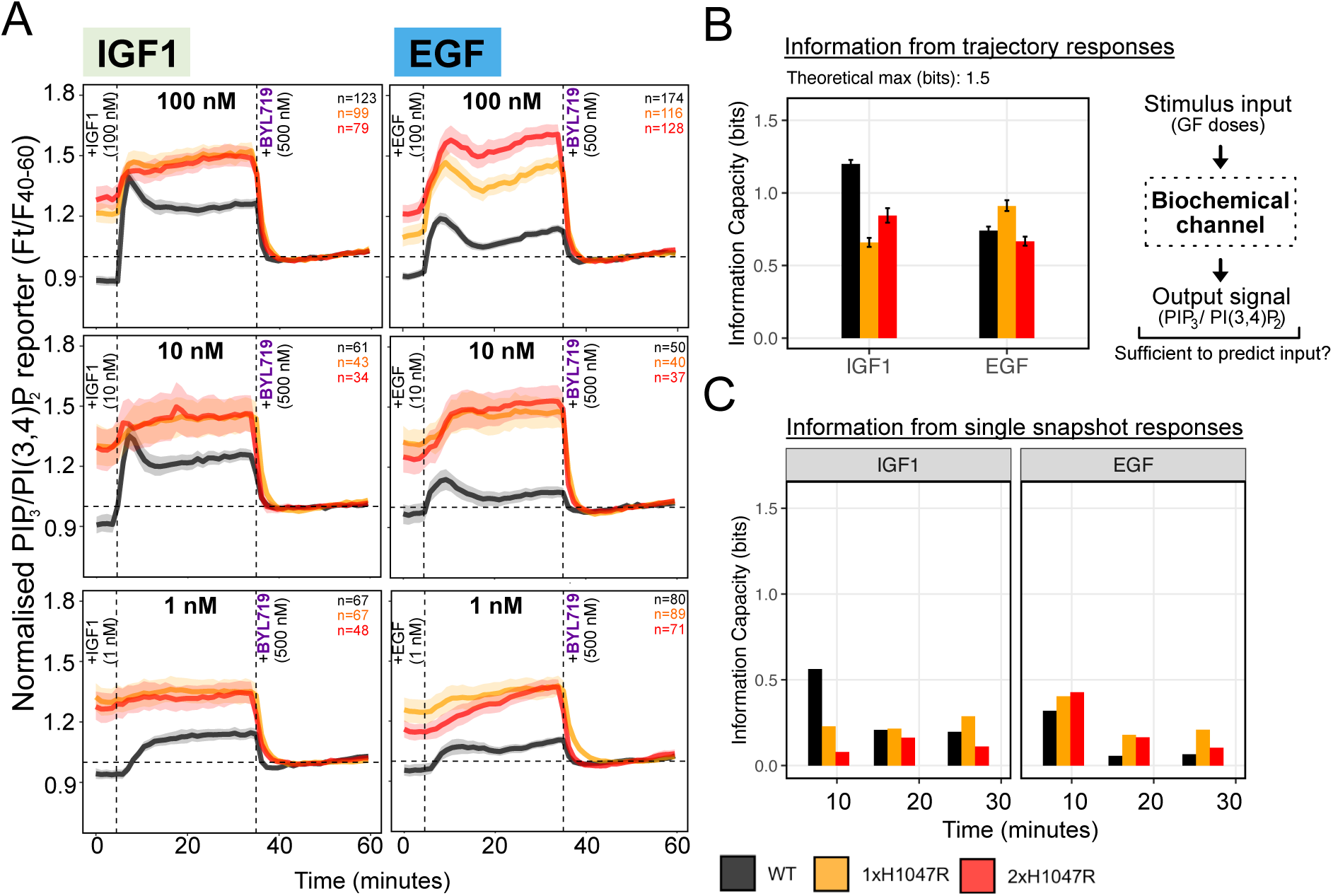
Oncogenic *PIK3CA^H1047R^* reduces the information capacity in the PIP_3_/PI(3,4)P_2_ dynamics for IGF1 but not EGF. **(A)** Total internal reflection (TIRF) microscopy measurements of IGF1- and EGF-induced PIP_3_/PI(3,4)P_2_ kinetics in live HeLa cells with endogenous, dose-controlled expression of *PIK3CA^H1047R^*(see also Fig. S2). Note that the wild-type traces correspond to those shown in Fig.1, shown separately across two figures for clarity. The cells were serum-starved for 3 h prior to stimulation with the indicated growth factors, followed by PI3Kα-selective inhibition with 500 nM BYL719. Measurements were obtained every 70 sec for a total of 60 min. The traces represent the mean PH_AKT2_ reporter fold-change relative to the median signal for the time window 35-60 min, used here to capture the baseline signaling elevation in *PIK3CA^H1047R^* mutant cells. The shaded areas represent bootstrapped 95% confidence intervals of the mean. The shading signifies the 95% confidence intervals of the mean. The number (n) of single cells for each genotype is indicated on the plots. For wild-type (WT) HeLa cells, two independent CRISPR/Cas9 clones were used, with and without silent mutations. The data are from 2 independent WT, 2 independent 1xH1047R and 3 independent 2xH1047R CRISPR/Cas9 clones. The data are from the following number (n) of independent experiments: n=6 for 100 nM IGF1; n=7 for 100 nM EGF; n=2 for 10 nM IGF1 and 10 nM EGF; n=3 for 1 nM IGF1; n=4 for 1 nM EGF. **(B)** Median information capacity in bits (log2) for IGF1 and EGF calculated from the trajectory responses in A. Capacity is a measure of the maximum amount of information that flows from the pathway input to its output. The theoretical maximum for 3 inputs (doses) is 1.5 bits if all the information is captured by the PIP_3_/PI(3,4)P_2_ dynamics. The error bars indicate interquartile range. **(C)** Median information capacity in bits (log2) calculated from snapshot measurements at the indicated time points from the datasets in A.

We found that HeLa cells with wild-type *PIK3CA* expression reached a relatively high mean information capacity of 1.2 bits for IGF1, suggesting that the majority of information about the dose of IGF1 is captured in the PIP_3_/PI(3,4)P_2_ response (**Fig. 2B**). Conversely, there was higher cellular uncertainty about the EGF doses based on the PIP_3_/PI(3,4)P_2_ trajectory alone, with *PIK3CA* wild-type HeLa cells reaching a mean information capacity of 0.75 bits. Notably, expression of the *PIK3CA^H1047R^* oncogene resulted in a substantial drop in information transfer downstream of IGF1, particularly in cells expressing a single copy of the mutation. Conversely, single-copy *PIK3CA^H1047R^*trended towards increased information capacity downstream of EGF (**Fig. 2B**).

Three conclusions can be drawn from these data. First, these results demonstrate that oncogenic *PIK3CA^H1047R^* can erode signaling fidelity in a growth factor-specific manner. Second, *PIK3CA^H1047R^* HeLa cells’ ability to distinguish between distinct doses of IGF1 on the basis of their PIP_3_/PI(3,4)P_2_ response degrades, reaching similar levels of information capacity as seen for EGF. Third, evaluation of temporal trajectories from the same cells and with high technical precision is key for accurate calculations of information transfer in signaling responses. As shown in **Fig. 2C**, information capacity calculations on snapshot measurements from the same data but without the temporal connection reveal erroneously low measures of PI3K signaling fidelity for both growth factors.

### *PIK3CA^H1047R^* corrupts the specificity of dynamic signal encoding

Further examination of the PIP_3_/PI(3,4)P_2_ trajectories in **Fig. 2A** suggested another key difference between wild-type and *PIK3CA^H1047R^*HeLa cells. Consistently, the EGF-induced PIP_3_/PI(3,4)P_2_ reporter response in mutant cells appeared amplified and largely indistinguishable from that of IGF1 in wild-type cells. This led us to hypothesize that *PIK3CA^H1047R^* expression may corrupt the cellular ability to resolve different growth factor inputs from one another. For this to have any significance, however, it would need to be reflected in the activity of key effectors downstream of PIP_3_/PI(3,4)P_2_ generation.

We therefore turned to live-cell imaging of a stably-expressed, high-fidelity FOXO-based kinase translocation reporter (KTR) (27), whose nucleocytoplasmic distribution provides a proxy measure for AKT activity (**Fig. S3A**) and is amenable to high-content-based, quantitative analyses (**Fig. S3B**). Compared to TIRF, widefield fluorescence imaging of the FOXO-based KTR response benefits from lower technical noise and allows capture of a much larger number of individual cells for robust information theoretic analyses across stimulations with different growth factors. We chose to compare IGF1, insulin, EGF and epigen due to their paired similarities at the level of activation of distinct RTKs (IGF1R/INSR vs EGFR). In *PIK3CA* wild-type cells, IGF1 and insulin elicited stronger and relatively similar FOXO-based KTR responses compared to EGF and epigen (**Fig. 3A, Fig. S3C,D**). Moreover, the temporal trajectories of IGF1/insulin remained highly distinct from those of EGF and epigen (**Fig. 3A, Fig. S3C,D**).

**Figure 3.**
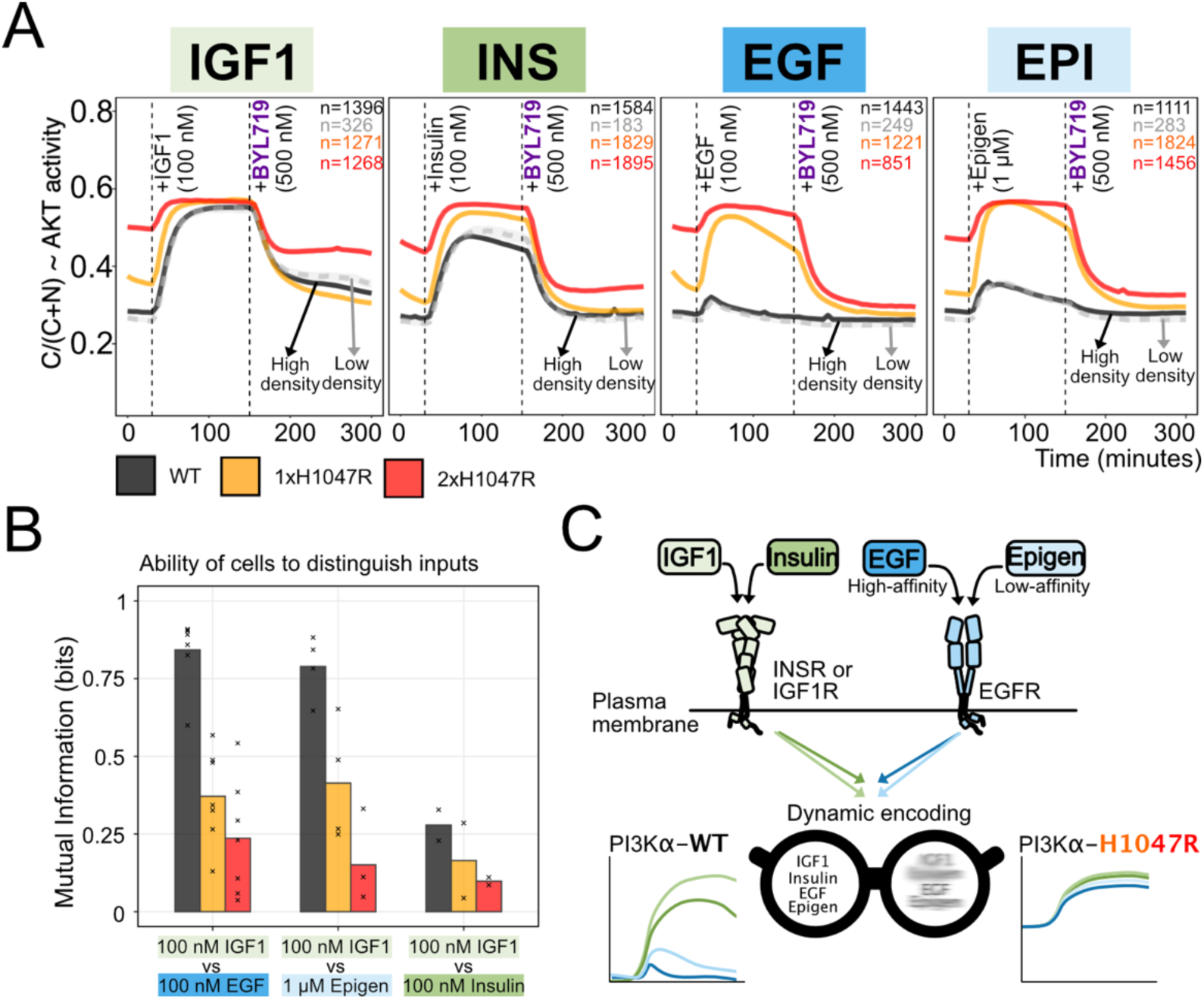
Oncogenic *PIK3CA^H1047R^*blurs the dynamic encoding of ligand identity. **(A)** Live-cell fluorescence-based measurements of a miniFOXO-based AKT kinase translocation reporter (KTR) (27), stably expressed in HeLa clones with the indicated *PIK3CA* genotypes. The total duration of the time course is 300 min, with measurements obtained every 6 min. For each time point, the traces correspond to the mean proportion of cytoplasmic KTR signal, with shaded areas representing bootstrapped 95% confidence intervals of the mean (note that these may be too small to be seen on the figure). C, cytoplasmic; N, nuclear. Single-cell numbers (n) are shown in the plots. For each condition, *PIK3CA* wild-type set were also seeded at low density to confirm intra-experimental consistency irrespective of cell crowding. The data are representative of a minimum of two independent experiments per condition, performed in two independent CRISPR/Cas9 clones per genotype. Plots from all independent experiments are shown in Fig. S3 and include control experiments with the 3xFS *PIK3CA* LOF mutant line. **(B)** Mutual information (MI) in bits (log2) for IGF1 *versus* each one of the indicated growth factors (EGF, epigen, insulin), calculated using the corresponding KTR trajectory responses (A) prior to inhibitor addition. MI values from individual experimental replicates are indicated as dots overlayed on barplots which correspond to the respective mean of each set of measurements. Note that IGF1 gave highly robust KTR dynamics, associated with relatively low single-cell noise as reflected in consistently high MI values. It was therefore chosen as control stimulus in all experimental replicates. **(C)** A graphic metaphor summarizing the biochemical signal blurring caused by oncogenic *PIK3CA^H1047R^*.

However, simply observing the average trajectories on their own is not enough to determine whether the FOXO-based KTR signaling dynamics are sufficiently distinct to allow individual growth factor inputs to be differentiated from one another. We therefore leveraged the entire set of single-cell trajectories to calculate the mutual information between IGF1 and every other growth factor. Mutual information takes into account the probabilistic and thus variable nature of individual growth factor responses. IGF1 was used as the control due to its highly robust single-cell KTR responses, both in terms of magnitude and temporal dynamics. Consistently, *PIK3CA^H1047R^* mutant cells exhibited an allele dose-dependent reduction in mutual information for all growth factors compared to IGF1 (**Fig. 3B**; note that mutual information is measured in bits, i.e., log2 scale). This drop was most pronounced for EGF and epigen, in line with the notion of selective amplification of the PI3K/AKT response downstream of EGFR seen in the mutant context. Consequently, the FOXO-based KTR response in mutant cells was no longer sufficiently distinct to resolve different growth factor inputs from one another. We therefore conclude that *PIK3CA^H1047R^* corrupts the dynamic encoding of signal identity, giving rise to cells with “blurred biochemical vision” (**Fig. 3C**).

### *PIK3CA^H1047R^* amplifies EGF signaling in cycling cells in 3D culture

A limitation of the approaches presented so far is reliance on exogenous reporters for evaluation of signaling responses. Moreover, TIRF-based measurements of the PIP_3_/PI(3,4)P_2_ reporter response are incompatible with joint tracking of the miniFOXO KTR reporter, limiting analyses to one response at a time. Finally, two-dimensional cell culture models do not capture the biological heterogeneity and additional complexity of three-dimensional (3D) culture systems. We therefore developed an orthogonal approach for single-cell-based signaling measurements in more complex culture settings, whilst retaining the ability to perform temporal perturbation experiments at scale.

Specifically, we adapted a recently published highly-multiplexed, mass cytometry workflow (28) for use with a new method that we developed for scalable generation of scaffold-free spheroids, including fixation for preservation of signaling responses and subsequent non-enzymatic single-cell dissociation (**Figure 4A**). We turned to mass cytometry given its versatility, compatibility with cell state-dependent gating and ability to multiplex up to 126 distinct conditions, with gains in sensitivity and technical robustness.

**Figure 4.**
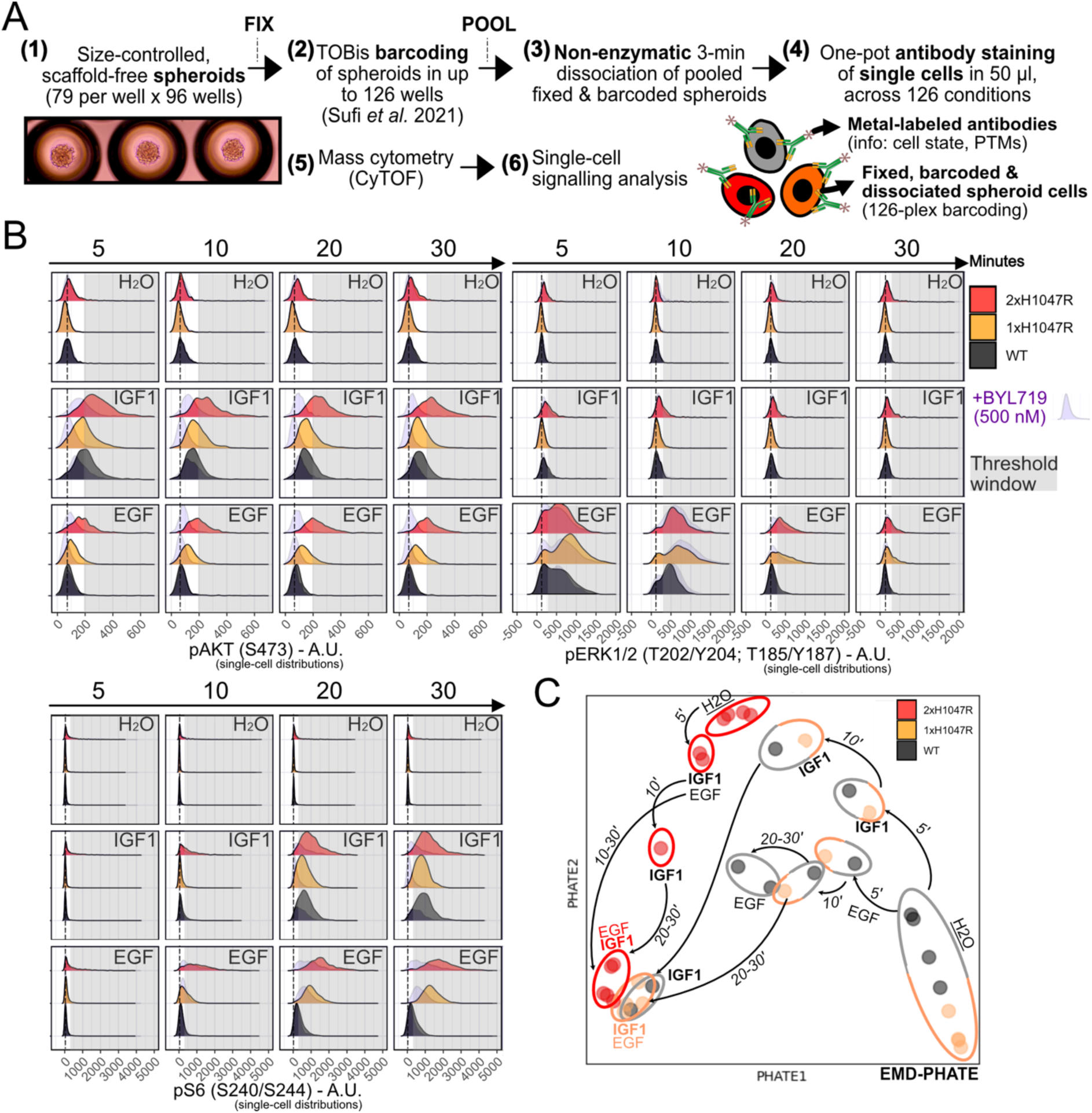
*PIK3CA^H1047R^*amplifies EGF-dependent signaling in a time- and allele dose-dependent manner. **(A)** Overview of the multiplexed mass cytometry workflow for profiling of single-cell signaling markers in scaffold-free spheroid models. Following fixation and thiol-reactive organoid barcoding in situ (83,28), up to 126 conditions are combined into a single sample for non-enzymatic single dissociation which ensures preservation of antibody epitopes, including post-translational modifications (PTMs). Subsequent staining with experimentally validated, metal-conjugated antibodies captures information about cell cycle state (e.g., cycling, non-cycling, apoptotic) as well as signaling state. **(B)** Mass cytometry (CyTOF) data from cycling, non-apoptotic HeLa spheroid cells with endogenous expression of wild-type (WT) *PIK3CA*, or one (1xH1047R) or two (2xH1047R) copies of the oncogenic *PIK3CA^H1047R^*. The spheroids were serum-starved for 4 h prior to stimulation with 100 nM EGF or IGF1, with and without 500 nM BYL719 (alpelisib; PI3Kα-specific inhibitor) as a control for signal specificity. Note that BYL719 was added at the same time as the growth factor, not as pre-treatment. The stippled line indicates the position of the peak in wild-type spheroids treated with vehicle (H_2_O). The grey shading highlights the response region that is not accessible to *PIK3CA* wild-type cells in the absence of stimulation. **(C)** Earth mover’s distance (EMD)-PHATE embedding of the signaling trajectories observed in the indicated HeLa cell genotypes. Single-cell distributions for the following signaling markers were used for EMD-PHATE processing (see also Fig. S5): pAKT^S473^, pERK1/2^T202/Y204; T185/Y187^, pNDRG1^T346^, pS6^S240/S244^, pSMAD2/3^S465/S467; S423/S425^.

Experiments with saturating doses of IGF1 and EGF revealed that robust growth factor signaling responses in HeLa spheroid cells were restricted to a cell cycling and non-apoptotic cell state (i.e., pRB Ser807/811-positive and cleaved Caspase3 D175-negative cells; **Fig. S4**). This aligns with recent findings of a multimodal, cellular state-conditioned sensitivity to growth factor stimulation in the human 184A1 breast epithelial cell line (29). Remarkably however, even when gating on the pRB^+^/cCASP3^-^ cell state, a comparison of the single-cell response distribution shifts relative to control treatments showed discernible, growth factor-specific temporal responses for phosphorylated AKT^S473^ (pAKT^S473^), ERK1/2^T202/Y204; T185/Y187^ and S6^S240/S244^ ribosomal protein (**Fig. 4B**). For example, in wild-type cells, the single-cell distribution shift for AKT phosphorylation was strongest upon stimulation with 100 nM IGF1 stimulation and peaked after 5-10 min. The same was true for ERK1/2 phosphorylation in response to EGF. Further downstream, a positive distribution shift for S6 phosphorylation (pS6^S240/S244^) followed with a delay relative to pAKT^S473^ (**Fig. 4B**), consistent with prior studies of bulk responses.

Next, to capture the temporal, signaling transitions present in this multidimensional dataset, we turned to PHATE (potential of heat diffusion for affinity-based transition embedding). PHATE produces a non-linear, low-dimensional embedding that preserves both local and global structure in the data (30). We first calculated an earth mover’s distance (EMD) score for each response distribution relative to untreated wild-type control cells; this score provides a concise measure of how different a single-cell distribution for a given signaling marker is relative to another (the corresponding wild-type control distribution in this case). PHATE was then applied to the EMD scores of all measured signaling markers (pAKT^S473^, pERK1/2^T202/Y204; T185/Y187^, pNDRG1^T346^, pS6^S240/S244^, pSMAD2/3^S465/S467; S423/S425^). This revealed a time-dependent convergence in signaling space of IGF1 and EGF responses for *PIK3CA^H1047R^* mutant cells (**Fig. 4C**), consistent with the “blurring” concept shown in **Fig. 3C**. This analysis also confirmed a clear distinction between HeLa cells with one *versus* two endogenous copies of *PIK3CA^H1047R^*, with the latter featuring higher baseline levels of the mTORC2 activation marker pNDRG1^T346^ (**Fig. S5A,B**).

In line with our 2D live-cell studies of PI3K/AKT signaling dynamics (**Fig. 1**, **Fig. 2**), we also observed that both single and two-copy *PIK3CA^H1047R^* mutant cells exhibited an amplified response to EGF stimulation (**Fig. 4B, 4C, S5A**). This resulted in stronger and more sustained responses both at the level of pAKT^S473^ and pERK1/2^T202/Y204; T185/Y187^. Relative to wild-type controls, the signaling responses in both single- and double-copy *PIK3CA^H1047R^* mutant HeLa cells also exhibited increased single-cell variability as a function of time and growth factor stimulation, most notably for pAKT^S473^ (**Figs. 4B**). This was reproduced with independent clones (**Fig. S5B**), and in additional dose-response, time course experiments using 1 nM, 10 nM, and 100 nM IGF1 or EGF (**Fig. S6A,B**). We therefore conclude that oncogenic *PIK3CA^H1047R^* does not simply shift the PI3K/AKT signaling response to a higher mean but also acts to enhance signaling heterogeneity.

### Corrupted signal transfer in *PIK3CA^H1047R^* mutant cell models translates into increased phenotypic heterogeneity in the context of EGF sensitization

We next set out to test whether the observed signal corruption in *PIK3CA^H1047R^* mutant cells translates into altered transcriptional and phenotypic responses. First, enhanced EGF signaling through AKT and ERK should lead to an amplification of EGF-specific transcriptional responses. These are known to be sensitive to the relative amplitude and duration of upstream signals such as ERK activation (31,32). Consistent with this prediction, we observed increased and more sustained mRNA expression of known EGF-dependent immediately early and delayed early genes in *PIK3CA^H1047R^*spheroids stimulated with EGF (**Fig. 5A**). This was specific to *PIK3CA^H1047R^*, as simply combining saturating concentrations of IGF1 and EGF to elicit strong activation of both AKT and ERK was not sufficient to amplify the transcriptional response in *PIK3CA* wild-type cells. Consistent with EGF’s known role as an epithelial-mesenchymal transition (EMT)-inducing factor (33,34), we also observed increased expression of the EMT-associated transcription factor *SNAIL (SNAI1)* in bulk *PIK3CA^H1047R^*HeLa spheroids (**Fig. 5A**). The apparent allele dose-dependent pattern of these responses was non-linear, however, with single-copy *PIK3CA^H1047R^*mutant cells exhibiting the strongest relative induction of EGF-dependent transcripts. These transcripts have also previously been associated with epithelial-mesenchymal transitions downstream of diverse inputs (33).

**Figure 5.**
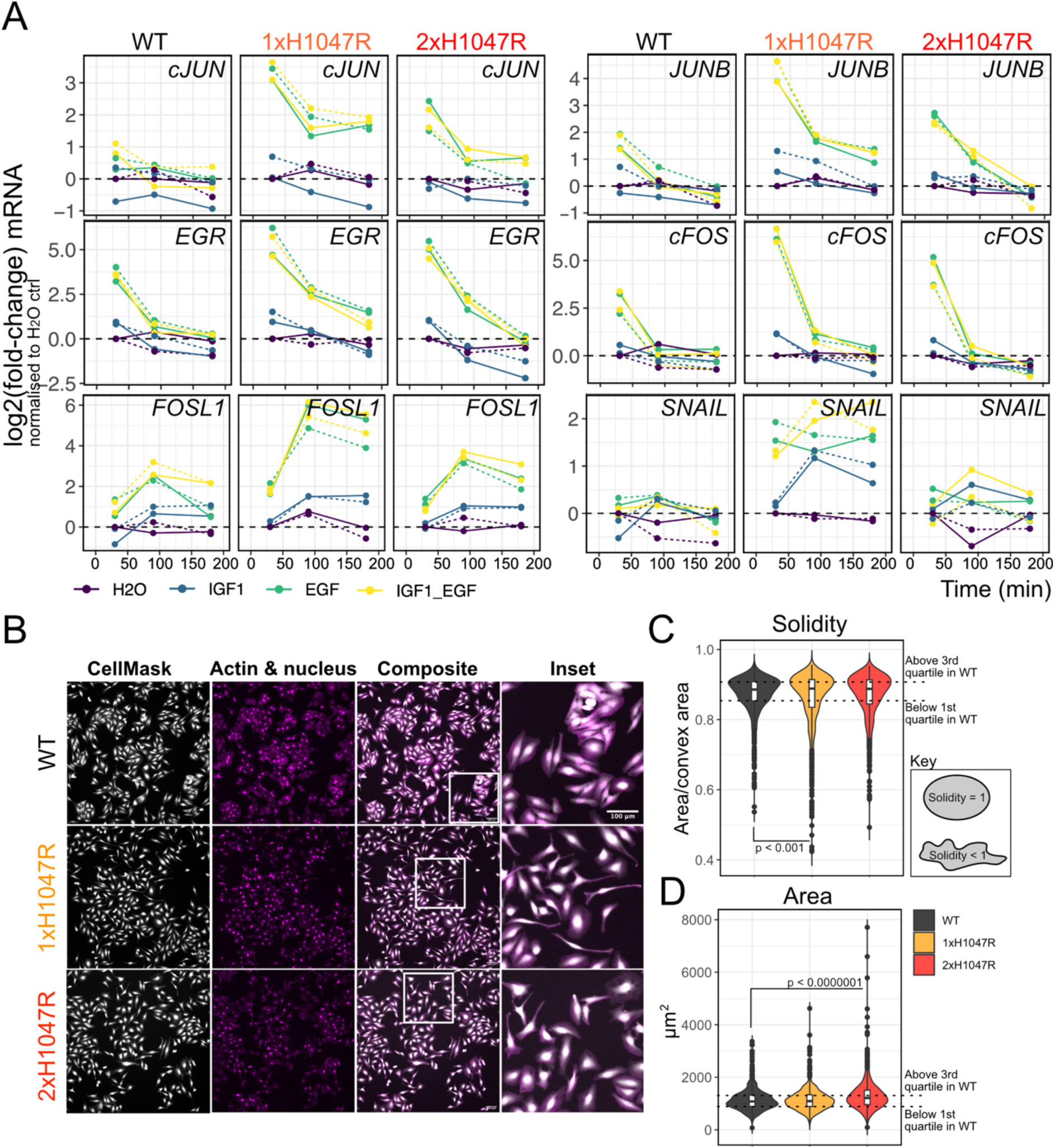
*PIK3CA^H1047R^* amplifies an EGF-driven transcriptional signature and increases phenotypic diversity in an allele dose-dependent manner. **(A)** Bulk transcriptional profiling of EGF-dependent immediate early and delayed early gene expression in HeLa spheroids with endogenous expression of either wild-type (WT) *PIK3CA*, one or two copies of *PIK3CA^H1047R^* (1-2xH1047R). Expression values are relative and represented as log2 fold-changes, normalized internally to each genotype’s control (H_2_O) response after 30 min of stimulation. All data were normalized to the expression values of *TBP* (housekeeping gene). The data are representative of two independent experiments (indicated with solid and stippled lines) with one CRISPR-derived clone per genotype. IGF1 and EGF were used at 100 nM, either alone or in combination as indicated. Note the log2 scale of the y-axis. **(B)** Representative fluorescence images of HeLa cells with the indicated genotypes during normal maintenance culture. The cells express a nuclear mCherry marker and were further stained with CellMaskBlue and Phalloidin to demarcate their cytoplasm and actin cytoskeleton, respectively. The cells are representative of three technical replicates and one CRISPR/Cas9 clone per genotypes (see also Fig. S6A for brightfield images of additional HeLa clones for each genotype). The cytoplasmic images from all replicates were used for deep learning-based segmentation (80), followed by quantification of cell shape solidity (**C**) and area (**D**). The scale bar in (B) corresponds to 100 µm. The p-values in **(C)** and **(D)** were calculated according to a one-way ANOVA with Tukey’s Honest Significant Difference to correct for multiple comparisons.

Second, the increased variability in signaling responses in *PIK3CA^H1047R^* HeLa cells would be expected to cause increased phenotypic heterogeneity. To evaluate this, we visualized cell appearance in standard 2D culture (**Fig. 5B**). Whereas wild-type HeLa cells grew as epithelial-like cell clusters, single- and double-copy *PIK3CA^H1047R^* cells were more dispersed and exhibited a higher proportion of cells with irregular, mesenchymal-like morphologies (**Fig. 5B-D, Fig. S7A)**. The mesenchymal shapes were most pronounced in single-copy *PIK3CA^H1047R^* mutant cells (**Fig. 5B,C**), in line with their higher expression of *SNAIL* upon EGF stimulation (**Fig. 5A**). Conversely, a higher proportion of the double-copy *PIK3CA^H1047R^* cells exhibited large, flattened morphologies (**Fig. 5B,D**).

The phenotypic heterogeneity observed in these HeLa cell models with endogenous, allele dose-dependent *PIK3CA^H1047R^*expression bore notable resemblance to the only other available model system of this kind – an allelic series of non-transformed, human iPSCs with heterozygous and homozygous expression of *PIK3CA^H1047R^*. Specifically, homozygous *PIK3CA^H1047R^* iPSC cultures were previously shown to exhibit coexisting epithelial and mesenchymal-like cellular morphologies (24) (Fig. S7B). We hypothesized that this phenotypic heterogeneity reflects corrupted signal transfer in homozygous *PIK3CA^H1047R^* iPSCs, including amplification of EGF-dependent responses and increased signaling heterogeneity as observed in the HeLa cervical cancer cell model. To test this, we applied our mass cytometry-based, single-cell signaling pipeline (**Fig. 4A**) on 3D-cultured, IGF1- or EGF-stimulated iPSCs from the aforementioned allelic series. As expected, both heterozygous and homozygous *PIK3CA^H1047R^* iPSCs had higher baseline phosphorylation of AKT (pAKT^S473^) relative to wild-type cells, and this increased further upon IGF1 stimulation (**Fig. 6A,B**). However, only homozygous *PIK3CA^H1047R^* iPSCs showed an amplified EGF response both at the level of AKT and ERK phosphorylation. Importantly, this was accompanied by a notable increase in the heterogeneity of the underlying *PIK3CA^H1047R/H1047R^*single-cell responses (**Fig. 6A,B**), revealing a conserved signaling phenotype that had remained inaccessible to conventional workflows based on bulk signaling measurements.

**Figure 6.**
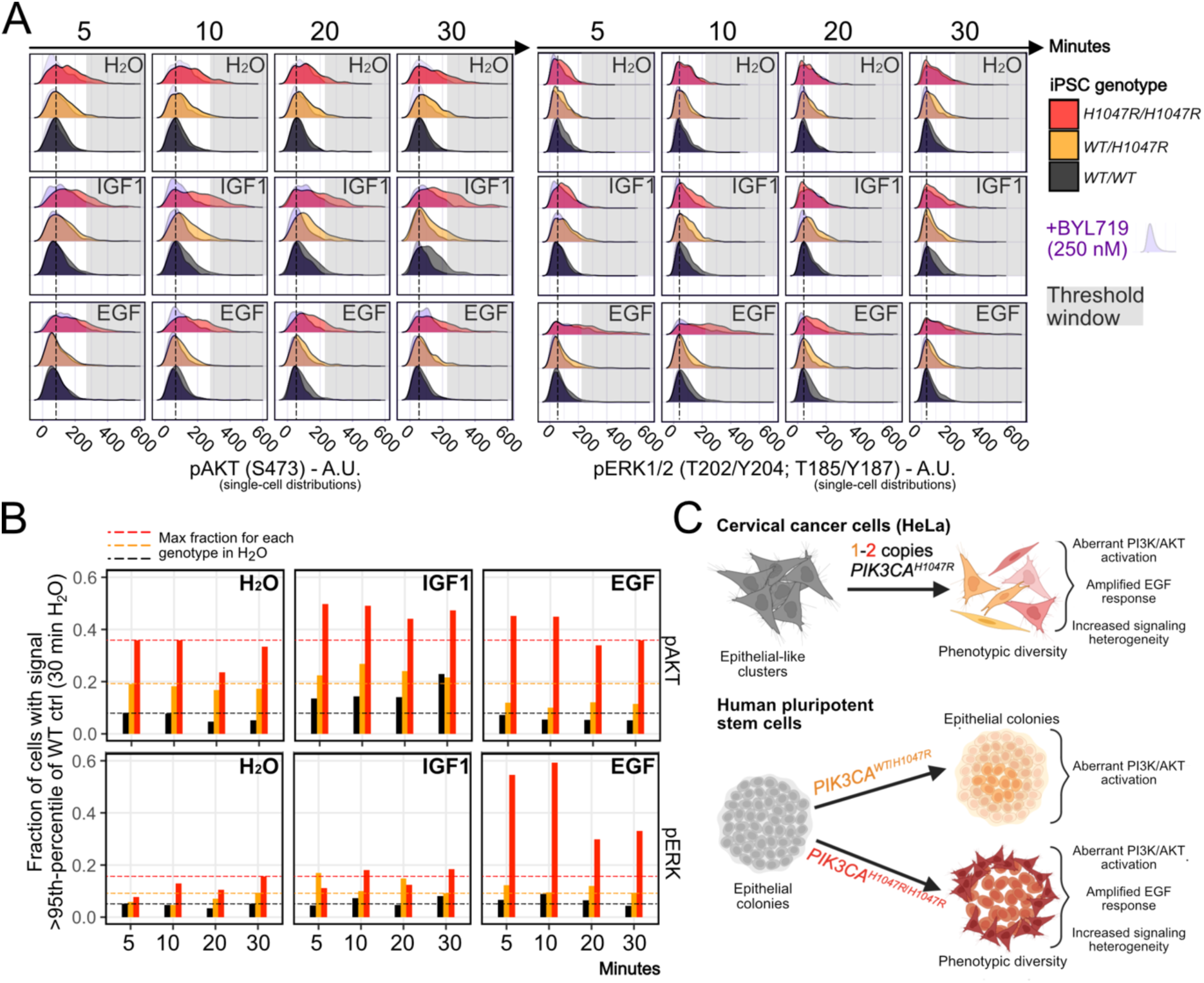
Corrupted signal transfer and EGF response amplification is conserved in homozygous *PIK3CA^H1047R^*iPSCs. **(A)** Mass cytometry data from cycling (pRB^+^) iPSC spheroid cells with wild-type (WT) *PIK3CA*, heterozygous or homozygous *PIK3CA^H1047R^*expression. The spheroids were serum-starved for 4 h prior to stimulation with 100 nM EGF or IGF1, with and without 250 nM BYL719 (alpelisib; PI3Kα-specific inhibitor) as a control for signal specificity. Note that BYL719 was added at the same time as the growth factor, not as pre-treatment. The stippled line indicates the position of the peak in wild-type spheroids treated with vehicle (H_2_O). The grey shading highlights the response region that is not accessible to *PIK3CA* wild-type cells in the absence of stimulation. **(B)** Thresholding of the data in (A) to quantify the percentage of cells within each condition with a pAKT or pERK signal above the corresponding 95^th^ percentile of vehicle (H_2_O)-treated WT iPSCs at 30 min. The stippled lines indicate the maximum fraction of cells within this threshold for each genotype prior to growth factor stimulation. The data are from a single experiment with one iPSC clone and n > 260 single cells per genotype. **(C)** Graphical summary of the key observations of the impact of *PIK3CA^H1047R^* expression in HeLa and iPSC cells.

In conclusion, compromised – or corrupted – signal transfer downstream of *PIK3CA^H1047R^* translates into increased signaling and phenotypic heterogeneity in the context of a selective amplification of EGF responses (**Fig. 6C**).

## Discussion

Despite tremendous advances in understanding of the core topology of the PI3K/AKT pathway over the last 30 years, the quantitative mechanisms of signal-specific information transfer in this pathway have remained elusive due to technical and analytical limitations (7,8). In this work, we present a suite of optimized single-cell-based, kinetic workflows for systematic mapping of quantitative signaling specificity in PI3K/AKT pathway activation. Supported by information theoretic analyses, we have shown that endogenous expression of the *PIK3CA^H1047R^* cancer hotspot variant results in quantitative blurring of growth factor-specific information transfer, amplification of EGF-induced responses and increased phenotypic heterogeneity in an allele dose-dependent manner. We note that an early study of bulk PI3K signaling responses in non-transformed breast epithelial cells also observed sensitization to EGF in the presence of either *PIK3CA^H1047R^* or the helical domain hotspot variant *PIK3CA^E545K^* (35). Our quantitative framework now allows these results to be contextualized into a coherent model of growth factor-specific mechanisms of action of oncogenic *PIK3CA*.

We propose a model in which oncogenic *PIK3CA^H1047R^* is not a simple ON switch of the PI3K/AKT pathway but acts as a context-dependent signal modifier, determined by the principles of quantitative biochemistry (8,36). Accordingly, the most parsimonious explanation for the selective amplification of EGF-dependent responses by *PIK3CA^H1047R^* is the ability of this mutation to increase p110α residency times at lipid membranes (37,38), alongside the lack of high-affinity phospho-Tyrosine (pTyr) binding sites for the regulatory p85 subunit on EGFR and its associated adaptor proteins (39). This would make the interaction between p110α and RAS essential for efficient PI3K-dependent signal transduction downstream of the EGFR (40–43), which would explain two key observations in our data: 1) the substantial increase in EGF-induced ERK phosphorylation in cell models expressing only mutant *PIK3CA^H1047R^*; 2) the complete suppression of AKT KTR responses following PI3Kα-selective inhibition in the context of EGF but not IGF1/insulin stimulation (Figs. 3 and S3). It is clear both from our PIP_3_/PI(3,4)P_2_ and AKT/FOXO (KTR) trajectories that IGF1 (and insulin) signals through both PI3Kα and an additional class IA PI3K isoform. This is unsurprising given the known ability of p85 to bind efficiently to pTyr (pYxxM) sites on IGF1R and insulin receptor substrate (IRS) proteins irrespective of the catalytic p110 subunit (44,45). It also explains the convergence of IGF1/insulin-induced AKT/FOXO (KTR) trajectories in *PIK3CA* loss-of-function and *PIK3CA^H1047R^* cells following PI3Kα-selective inhibition (Fig. S3). The only cell line in this work in which PI3Kα appeared dispensable for EGF-induced PI3K signaling was A549, a lung adenocarcinoma cell line with amplified EGFR (46). This aligns with the above biochemical considerations; a higher concentration of EGFR would allow for more successful engagement of low-affinity interactors such as p85, thus making PI3K activation less dependent on RAS binding via p110α. This follows from the fact that p110β, which is the other main catalytic p110 isoform in the non-hematopoietic cell lineages used here, does not have the capability of direct interaction with RAS (47).

The above model is nevertheless a simplification because it does not account for another salient property of oncogenic *PIK3CA,* revealed by our quantitative single-cell measurements of PI3K/AKT signaling dynamics. Thus, we consistently observed an increase in signaling and phenotypic heterogeneity downstream of allele dose-dependent *PIK3CA^H1047R^* expression. It is interesting that an increased heterogeneity in PI3K pathway activation was also noted in an early study of *PIK3CA^H1047R^* overexpression in breast epithelial cells (48). The consequences of this heterogeneity are two-fold. First, it increases the uncertainty in the ability to predict the outputs of oncogenic PI3K/AKT pathway activation, i.e. the outputs are probabilistic rather than deterministic. This calls for increased attention to single-cell PI3K signaling responses in the ongoing evaluation of the many PI3K/AKT pathway inhibitors entering preclinical and clinical use (4,49,50). Second, such heterogeneity endows cells with the ability to sample multiple phenotypic states, or attractors (51), as shown by the emergence of co-existing cellular phenotypes in otherwise isogenic cells with *PIK3CA^H1047R^* expression. This may offer a mechanistic underpinning for the remarkable phenotypic heterogeneity found in *PIK3CA*-driven breast cancer models (52–54) as well as benign but highly debilitating human PROS disorders (5). It is likely that the observed increase in single-cell “noise” endows the population of mutant *PIK3CA^H1047R^* cells with a selective advantage in the face of unpredictable and rapidly changing environments as shown in other systems (55).

Finally, our quantitative PI3K signaling framework also makes possible the prospective development and application of pharmacological approaches to tune pathological PI3K signaling responses back to normal, for example through allosteric modulation of receptor-specific coupling mechanisms. This has also been suggested for RAS/MAPK signaling (56), in light of recent findings that oncogenic mutations in this pathway also remain dependent on upstream growth factor inputs yet fail to transmit these reliably (9). Given the likely dependence on a direct PI3Kα-RAS interactions for the EGF response amplification in *PIK3CA^H1047R^* cells, the recently released PI3Kα-RAS breaker (57) may be an excellent candidate for testing of quantitative, growth factor-specific PI3K signaling dynamics as a pharmacological target.

## MATERIALS AND METHODS

Where referenced, the Open Science Framework (OSF) project containing all source datasets underpinning this work can be accessed via doi: 10.17605/OSF.IO/4F69N.

### Immortalized cell culture

HeLa cells and mouse embryonic fibroblasts (MEFs) were cultured in complete medium consisting of DMEM (with 4 mM L-Glutamine and 1 mM sodium pyruvate; Thermo Fisher Scientific #41966-029) supplemented with another 2 mM of L-Glutamine (Sigma #G7513) and 10 % fetal bovine serum (FBS; Pan-Biotech #P30-8500).

Lung adenocarcinoma A549 cells were cultured in complete medium consisting of RPMI-1640 with GlutaMax and Sodium Bicarbonate (#61870-036, Thermo Fisher Scientific), supplemented with 1 mM of Sodium Pyruvate (#11360-039, Thermo Fisher Scientific) and 10 % FBS.

Cells were cultured in T25 flasks (Corning or TPP) and passaged every two-to-three days when 80-90% confluent. Briefly, the spent medium was removed and the cells washed with 5 ml DPBS (Sigma #RNBH8966 or Thermo Fisher Scientific # 14190-094). Following removal of the wash, the cells were incubated at 37°C in 0.75 ml TrypLE™ Express Enzyme (Thermo Fisher Scientific #12605028 or #1260421) for 6-8 min until dissociated. The cells were resuspended in complete medium and distributed to new flasks at appropriate ratios.

### Human induced pluripotent stem cell culture

The male human iPSCs used in this work were derived from the WTC11 line, following CRISPR/Cas9 engineering for endogenous expression of *PIK3CA^H1047R^*as described previously (24). The cells were maintained in Essential 8 Flex Medium (Thermo Fisher Scientific #A2858501) on plates coated with 10 µg/cm^2^ Cultrex Stem Cell Qualified Reduced Growth Factor Basement Membrane (R&D Systems #3434-010-02). Cells were cluster-passaged every 3-4 days with 0.5 mM EDTA and seeded into medium supplemented with 10 µM Y-27632 dihydrochloride (Bio-Techne #1254/10) for the first 24 h. For details of the spheroid set-up, see dx.doi.org/10.17504/protocols.io.3byl4bnrrvo5/v1. The cells were single-cell dissociated with StemPro Accutase (Thermo Fisher Scientific # A1110501) and seeded at 1000 cells/spheroids in 200 µl Essential 8 Flex supplemented with 10 µM Y-27632 dihydrochloride. The following day, the medium was replenished without Y-27632. Spheroids were processed for experimental perturbations two days following formation.

### Cell line quality control

All cell lines were cultured in the absence of antibiotics except if processed for selection post-engineering as indicated. Cells were routinely tested negative for mycoplasma and genotyped by Sanger sequencing (knock-in lines) or immunoblotting (knock-out lines) to confirm the correct identity prior to experimental use.

### CRISPR/Cas9 gene editing of *PIK3CA* exon 21 in HeLa cells

Low-passage (P5) HeLa cells were used for CRISPR/Cas9 engineering for knock-in of the *PIK3CA* H1047R variant (c.CAT>c.CGT) using a modified version of the protocols described in Refs. (24,58). Briefly, a total of 200,000 cells were targeted with a total of 200 pmol single-stranded oligodeoxynucleotides (ssODNs) introducing either the targeting mutations along with silent mutations or silent mutations without the targeting mutation: *HDR001 ssODN (targeting mutation and silent mutations): 5’-TAGCCTTAGATAAAACTGAGCAAGAGGCTTTGGAGTATTTCATGAAACAAATGAACGACGCACGTCATGGTGGCTGGACAACAAAAATGGATTGGATCTTCCACACAATTAAACAGCA TGCATTGAACTGAAAAGATAACTGAGAAAATG-3’*

*HDR002 ssODN (silent mutation only): 5’-TAGCCTTAGATAAAACTGAGCAAGAGGCTTTGGAGTATTTCATGAAACAAATGAACGACGCACATCATGGTGGCTGGACAACAAAAATGGATTGGATCTTCCACACAATTAAACAGCATGCATTGAACTGAAAAGATAACTGAGAAAATG-3’*

Three different mixtures of ssODNs (HDR001 alone, HDR002 alone, 1:1 mixture HDR001:HDR002) were set up to ensure generation of a dose-controlled allelic series for *PIK3CA^H1047R^*. Targeting was performed using recombinant ribonucleotide proteins (RNPs) at ratio 1:1.2 (Cas9:sgRNA; 4 µM:4.8 µM). The synthetic sgRNA (5’-AUGAAUGAUGCACAUCAUGG-3’) was obtained from Synthego (modified for extra stability). The high-fidelity Alt-R™ S.p. Cas9 Nuclease V3 (IDT # 1081061) was used to limit off-targeting risk. Cells were targeted by nucleofection using the SE Cell Line 4D-Nucleofector™ X Kit S (Lonza #V4XC-1032). Cells were allowed to recover from nucleofection before sib-selection-based subcloning to isolate pure clonal cultures. To aid recovery, conditioned medium (1:1 mixture with fresh medium) was used for 7 days during subcloning.

An initial screen for correct genotypes was performed using DNA extracted with QuickExtract (Cambridge Bioscience # QE0905T) and subjected to PCR amplification and Sanger sequencing with primers: 5’-CAGCATGCCAATCTCTTCAT-3’ (forward), 5’-ATGCTGTTCATGGATTGTGC-3’. As HeLa cells are triploid on average, genotypes were called following deconvolution with Synthego’s ICE tool (59). Putative pure wild-type (or silent mutation only) and *PIK3CA^H1047R^* clones were expanded and subjected to final validation by next-generation sequencing using MiSeq, with Illumina adaptor-appending primers:

*5’-TCGTCGGCAGCGTCAGATGTGTATAAGAGACAGATAAAACTGAGCAAGAGGCTTTGGA-3’ (forward);*

*5’-GTCTCGTGGGCTCGGAGATGTGTATAAGAGACAGATCGGTCTTTGCCTGCTGAG-3’ (reverse)*.

The MiSeq output was analyzed using CRISPResso2 (60). All raw and analyzed files are deposited on the accompanying OSF project site (doi: 10.17605/OSF.IO/4F69N, component: *MiSequencing_CRISPR_clone_validation*).

Note that HeLa cells are nominally triploid. All knock-in lines with one or two copies of *PIK3CA^H1047R^* therefore harbor two or one allele(s), respectively, with a C-terminal frameshift equivalent to those in the loss-of-function 3xFS clone (**Fig. S2**). This frameshift is too close to the stop codon of p110α to result in nonsense-mediated decay and instead causes a change of the last 20-30 C-terminal amino acids. This abolishes the critical p110α WIF motif required for membrane binding and catalytic function (38), effectively creating a loss-of-function knock-in that nevertheless remains expressed and thus does not carry the risk of altering the stoichiometry of p85 regulatory and p110 catalytic subunits in these cells. In separate QC tests, we also used a clone with 2xWT and 1xFS *PIK3CA* alleles and confirmed that its PIP_3_/PI(3,4)P_2_ reporter response by TIRF microscopy is identical to those in the 3xWT clones. Combined with data from the 3xFS clone, this provides confidence that the frameshift p110α allele(s) present alongside *PIK3CA^H1047R^* do(es) not impact the observed signaling responses.

### Exome sequencing

All CRISPR/Cas9-engineered HeLa clones used in this work were profiled by whole-exome sequencing at an early passage (13 or 14) and compared to the parental HeLa culture prior to editing (passage 4) and following another 10 passages (passage 14) in the absence of editing. This approach enabled rigorous evaluation of the extent of mutagenesis caused by the gene editing and single-cloning procedures relative to the expected baseline acquisition of mutations upon prolonged cell culture. High-quality DNA was extracted using the NucleoSpin Micro Kit XS (Takara #740901.5) and submitted to Novogene for exome library preparation and sequencing. Briefly, libraries were prepared with the Next Ultra DNA Library Prep Kit (NEB #E7370L) and enriched for exons using Agilent’s SureSelectXT Reagent Kit and Agilent SureSelect Human All ExonV6 (#G9611B). The final libraries were pooled and paired-end (150 bp) sequenced on a NovaSeq 6000 instrument, with 6G of raw data output per sample. Subsequent read processing was performed with the nf-core/sarek pipeline (v2.6.1), with alignment against the human genome (hg38) and Agilent’s reference .bed file corresponding to the SureSelect Human All Exon V6 60MB S07604514 design. Somatic variant calling was performed with Strelka2 (61) according to a tumor/normal pairs setup where CRISPR/Cas9-edited clones and the long-term passage parental cultures were assigned the “tumor” label and the low-passage parental culture assigned the “normal” label. Subsequent variant annotation was performed using both the snpEff (62) and VEP pipelines (63). The VEP pipeline, however, failed to capture the H1047R variants in the mutant lines correctly and its output was therefore deemed unreliable. Detailed scripts and multiQC reports for reproducing the all nf-core/sarek outputs have been made available on the OSF project site.

The snpEff-annotated variants with SomaticEVS filter = PASS were processed with GATK VariantsToTable and imported into R for identification of non-synonymous protein-coding variants that are common to a minimum of two samples when compared to the low-passage parental culture prior to CRISPR/Cas9 gene editing. Intersection plots and heatmaps were generated using the ComplexHeatmap R package (64). Clustering was performed according to Euclidean distance with the Ward.D2 method. Raw sequencing data and annotated R processing scripts are provided on the accompanying OSF project site (doi: 10.17605/OSF.IO/4F69N, component: *Exome_sequencing_processed_file_analysis*).

### Total mRNA sequencing

All CRISPR/Cas9-edited HeLa clones were processed for total mRNA sequencing at baseline to determine transcriptional similarities and differences across individual genotypes. Individual clones (passages 16-18) were collected at subconfluence following refeeding with fresh complete medium for 3 h. Following a single wash with DPBS, cells were snap-frozen and stored at −80°C until further processing. Following thawing on ice, total RNA was extracted using the Direct-zol RNA Miniprep Kit from ZymoResearch (#R2051), with final elution in 30 µl nuclease-free water. Samples were submitted to Novogene for library preparation (NEB Next® Ultra™ RNA Library Prep Kit) and paired-end (150bp) sequencing on a NovaSeq 6000 instrument. Note that the library preparation is strand-agnostic.

Raw read processing was performed with the Nextflow (version 20.07.1) nf-core RNAseq pipeline (v1.1) (65), with Spliced Transcripts Alignment to a Reference (STAR) (66) for read alignment to the human genome (Homo_sapiens.GRCh38.96.gtf) and featureCounts (67) for counting of mapped reads (multimapped reads were discarded). All subsequent data processing was performed in R, with differential gene expression analysis following the limma-voom method (68). Filtering of low gene expression counts was performed with the TCGAbiolinks package with quantile value 0.75 (chosen empirically based on the observed count distribution). Next, read count normalization was performed with the gene length-corrected trimmed mean of M-values (GeTMM) method (69). PCA was done using the PCAtools package. The mean-variance relationship was modelled with voom(), followed by linear modelling and computation of moderated t-statistics using the lmFit() and eBayes() functions in the limma package (68). Experimental replicate was included as a batch effect term in the model. The associated p-values for assessment of differential gene expression were adjusted for multiple comparisons with the Benjamini-Hochberg method at false-discovery rate (FDR) = 0.05 (70). Adjustments were performed with option = “separate”, comparing *PIK3CA^H1047R^*mutant clones against wild-type clones. No differentially expressed genes were identified across the different genotypes at baseline. All R processing scripts to replicate the analyses are provided on the accompanying OSF project site (doi: 10.17605/OSF.IO/4F69N, component: *RNAseq_processing*). The raw sequencing files are available via GEO under accession number: GSE251956.

### Sleeping Beauty transposon engineering of cells for expression of AKT kinase translocation reporter (KTR)

The Sleeping Beauty transposon-based and optimized miniFOXO kinase translocation reporter (KTR) developed by Gross *et al.* (27) was used to generate stable cell lines from the original CRISPR/Cas9-engineered HeLa cell clones. This was performed at two different sites in two independent sets of wild-type and mutant clones, with a time gap of one year. For further testing of reproducibility of the results irrespective of reporter expression levels, stable cell line generation was performed using two different molar ratios of transposon to transposase (the following molar units are for cells seeded in 12-well plates at a density of 50,000 cells/well; these units were scaled by a factor of 2 for cells seeded in 6-well plates at 100,000 cells/well). For a 1:1 molar ratio, engineering was performed with approximately 100 fmol of transposon and SB100X transposase-expressing plasmids (the plasmid maps are deposited on the OSF project site, with code names MB40 and MB43, respectively). For a 1:10 molar ratio and thus low reporter expression, engineering was performed with approximately 10 fmol transposon plasmid and 100 fmol transposase plasmid. Plasmid were delivered to cells using Fugene HD Transfection Reagent (Promega #E5912) at a 3:1 Fugene volume:DNA mass ratio for transfection complex formation in Opti-MEM I Reduced Serum Medium (Thermo Fisher Scientific #31985070). Puromycin (Sigma Aldrich #P9620 or #P4512-1MLX10) selection at 1 µg/ml was started 24-48 h after seeding, with replenishment of selection medium at least every second day. Stable cell lines were usually established and banked within 2 weeks of the initial transfection.

### Western blotting

A step-by-step Western blotting protocol is publicly available on protocols.io with the following doi: dx.doi.org/10.17504/protocols.io.4r4gv8w. Cells were lysed from 10-cm dishes with RIPA Lysis and Extraction buffer (Thermo Fisher Scientific #89900), and 10-15 µg of protein were loaded on 4-12% Bis-Tris Midi NuPage Protein Gels (Thermo Fisher Scientific) and separated at 120V for 2 h in MES running buffer. Protein transfer was performed with an iBlot2 system (Thermo Fisher Scientific) using program P3. All primary and secondary antibodies used are provided in Tables 1 & 2. Final signal detection was by enhanced chemiluminescence (ECL) with the Immobilon Forte Western HRP substrate from Sigma Aldrich (#WBLUF0500) or ECL Western Blotting Substrate from Promega (#W1015). Images were acquired on the Amersham ImageQuant 800 system with 5×5 binning. All raw Western blots have been deposited on the accompanying OSF project site (doi: 10.17605/OSF.IO/4F69N, component: *Western_blots_Fig.S2*).

### Small molecule reconstitution and usage

The following growth factors were obtained from Peprotech: human IGF1 (#100-11, lots: 022201-1, 092101-1, 041901-1), human EGF (#AF-100-15, lots: 0922AFC05, 0222AFC05, 0820AFC05), human Epigen (#100-51, lot: 0706386). Lyophilised stocks were reconstituted in sterile, molecular-grade, non-DEPC-treated water from Ambion (#9937), allowed to dissolve for 15-20 min at 4°C followed by aliquoting in PCR strip tubes and long-term (up to 1y) storage at −80°C. Aliquots were freeze-thawed maximum once to limit loss of potency.

Human insulin (10 mg/ml) was from Sigma (#91077C, lot: 21M018) and stored at 4°C.

BYL719 was obtained from SelleckChem (#S2814, lots: 03, 06) at 10 mM in DMSO. The stock solution was diluted to 1 mM in sterile DMSO, aliquoted in PCR strip tubes and stored at −80°C long-term (up to 2 years).

TGX221 was obtained from MedChemExpress (#HY-10114) and reconstituted in sterile DMSO at 10 mM DMSO, prior to long-term storage (up to 3 years) at −80°C.

1938 was synthesised by Key Organics or SAI Life Sciences, and is now available through CancerTools (#161068).

### Phosphoinositide reporter constructs

To minimize confounding effects on reporter performance arising from usage of different plasmid backbone, all PH domain derivatives were cloned into the same plasmid backbone construct (pNES-EGFP-C1 for wild-type PH domains; pNES-mCherry-C1 for mutant PH domains). The generation of each individual reporter is detailed below. Note that all PH domains will now be coupled to a nuclear export sequence (NES) and harbor an N-terminal fluorescent protein tag. All plasmids were verified by restriction enzyme digest and Sanger sequencing. Plasmid maps have been deposited on the accompanying OSF project site (doi: 10.17605/OSF.IO/4F69N; component *Other*).

#### ARNO

The pNES-EGFP-C1-PH-ARNO(I303E)x2 PIP_3_ reporter construct was a gift from Dr Gerry Hammond (University of Pittsburgh) and was generated as described in Ref. (18). In this construct, the PH domain is C-terminally tagged with an enhanced GFP (EGFP) which is itself preceded by a nuclear export sequence. To generate a tandem-dimer mutant version equivalent to R280A in the native PH domain of ARNO (Unitprot #P63034), a 996 bp gene fragment corresponding to the tandem-dimer PH-ARNO(I303E) domain with the mutated residues was synthesized as a gene fragment in a pUC vector by GeneWiz, including 5’ and 3’ HindIII and BamHI recognition sites, respectively. Next, five reactions each with 250 fmol of the construct carrying the mutant fragment or the original wild-type pNES-EGFP-C1-PH-ARNO(I303E)x2 construct were digested with 20 U each of BamHI-HF (NEB #R3136S) and HindIII-HF (NEB #R3104S), alongside 5 U of quick alkaline phosphatase (calf intestinal, NEB #M0525S), all in a 30 µl rCutSmart buffer (NEB) reaction. The digests were run at 37°C overnight (16 h), followed by heat inactivation at 80°C for 20 min. The digests were then un on a Tris acetate-EDTA agarose gel (1%), followed by gel purification of the pNES-EGFP-C1 destination vector and the mutant PH-ARNO(I303)x2 domain with compatible sticky ends, using the Monarch DNA gel extraction kit (NEB #T020S) according to the manufacturer’s instructions. The insert and the destination vector were ligated in a 10 µl reaction with 2X instant sticky-end ligase master mix (NEB #M0370S), using a 1:5 molar ratio of backbone-to-insert and otherwise following the manufacturer’s instructions. Next, 2 µl of the ligation reaction were heat-shock transformed into high-efficiency 5-alpha competent *E. coli* (NEB #C2987I), followed by conventional colony picking and bacterial culture expansion for subsequent plasmid DNA extraction with the Maxi Plus kit from Qiagen (#12964). Next, the EGFP tag in the new construct containing the mutant PH-ARNO(I303)x2 domain was replaced with an mCherry tag obtained from a pNES-mCherry-C1-TAPP1-cPHx3 construct (a gift from Dr Gerry Hammond; described in Ref. (18)). The restriction enzyme digest-based subcloning protocol used to generate this construct was as described above.

#### BTK

The BTK-PH domain was obtained from Addgene construct #51463 (a gift from Dr Tamas Balla). This construct was used for site-directed mutagenesis of a key arginine in the signature motif (FKK<u>R</u>L) of the BTK PH domain (20) using the following primers: 5’-CTTCAAGAAGgcCCTGTTTCTCTTG-3’ (forward); 5’-TTTAGAGGTGATGTTTTCTTTTTC-3’ (reverse). Site-directed mutagenesis was performed with the Q5® Site-Directed Mutagenesis Kit from New England Biolabs (#E0554S), using 0.5 µM of each primer and 0.2 ng/µl plasmid DNA in a 25-µl reaction. The thermocycling conditions were as follows: denaturation at 98°C for 30s; 25 cycles of 98°C for 10s, 56°C for 20s, 72°C for 2.5 min; final extension at 72 °C for 5 min. The PCR product was subsequently processed for KLD (kinase, ligase, DpnI) treatment as per the manufacturer’s instructions.

The wild-type and mutant versions of the BTK PH domain were PCR-amplified, including addition of 5’ and 3’ BamHI and HindIII restriction enzyme recognition sites, respectively. The following primers were used (with highlights to indicate the recognition sites): 5’-AGCAGAAGCTTCGATGGCCGCAGTGATTCTGG −3’ (forward); 5’-CCGGTGGATCCTCACCGGATTACGTTTTTGAGCTGG −3’ (reverse; note this primer also adds a stop codon). A two-step PCR amplification was performed using Platinum SuperFi DNA Polymerase (Thermo Fisher Scientific #12351-010) with the following thermocycling conditions: denaturation at 98 °C for 30s; 5 cycles of 98°C for 10s, 60°C for 10s, 72°C for 15s; 20 cycles of 98°C for 10s, 65°C for 10s, 72°C for 15s; final extension at 72°C for 5 min. The PCR products were gel-purified and processed for HindIII- and BamHI-based subcloning into the pNES-EGFP-C1 and pNES-mCherry-C1 backbones as described above for ARNO.

#### AKT2

The AKT2-PH domain was obtained from a plasmid encoding the full length AKT2 protein (a gift from Dr James Burchfield; described in Ref. (71)). This construct was used for site-directed mutagenesis of a key arginine in the signature motif (WRP<u>R</u>Y) of the AKT2 PH domain (20) using the following primers: 5’-CTGGAGGCCAgcGTACTTCCTG-3’ (forward); 5’-GTCTTGATGTATTCACCAC-3’ (reverse). Site-directed mutagenesis was performed as described for BTK except for use of 57°C as annealing temperature and 4 min of extension time in each cycle. The wild-type and mutant versions of the AKT2 PH domain were PCR-amplified, including addition of 5’ and 3’ BamHI and HindIII restriction enzyme recognition sites, respectively. The following primers were used (with highlights to indicate the recognition sites): 5’-AGCAGAAGCTTCGATGAATGAGGTGTCTGTCATC-3’ (forward); 5’-CCGGTGGATCCTCAGTTGGCGACCATCTGGA-3’ (reverse; note this primer also adds a stop codon). The procedure was as described for BTK above, with subsequent subcloning into the pNES-EGFP-C1 and pNES-mCherry-C1 backbones as described for ARNO.

### Live-cell total internal reflection fluorescence (TIRF) microscopy

A detailed protocol of how to prepare HeLa cells for live-cell microscopy by TIRF, including Matrigel coating of the dishes, cell seeding, transfection with phosphoinositide reporters and subsequent treatment has been made publicly available on protocols.io via the following doi: dx.doi.org/10.17504/protocols.io.kxygx37jkg8j/v1

The above protocol was also followed for experiments with MEFs and A549 with the following modifications. MEFs were seeded at a density of 2000 cells per well (0.35 cm^2^). A549 cells were seeded either at 2000 or 3000 cells per well and transfected either with 25 ng or 50 ng wild-type and mutant reporter constructs; these different conditions were tested due to the low transfection efficiency of these cells, however the final results did not differ and were thus pooled together. Note that 20 ng of a pUC19 (NEB #09052008) carrier plasmid was included in all transfection conditions with 25 ng of each reporter plasmid for a more even uptake of the latter.

Time-lapse TIRF images were obtained on a 3i Spinning Disk Confocal microscope fitted with a sCMOS Prime95B (Photometric) sensor for TIRF, with full temperature (37°C) and CO_2_ (5%) control throughout the acquisitions. A 100X 1.45 NA plan-apochromatic oil-immersion TIRF objective was used to deliver the laser illumination beam (488 nm or 561 nm; 40-50% power) at the critical angle for TIRF and for acquisition of the images by epifluorescence (200-300 msec exposure) using single bandpass filters (445/20 nm and 525/30 nm). Acquisition was performed in sequential mode, without binning, using Slidebook 6.0 and an acquisition rate of 70s.

Image analyses of total reporter intensities were performed with the Fiji open source image analysis package (72). The regions of interest (ROI) corresponding to the footprint of the individual cell across time points were defined using minimal intensity projection to select only pixels present across all time points, following prior background subtraction with the rolling ball method (radius = 500 pixels) and xy drift correction. Mean intensity levels for each reporter were measured within the ROI and exported for subsequent data processing in R. Final trajectory normalizations to the median signal of pre-stimulus or post-BYL719 time points were performed using the Time Course Inspector package *LOCnormTraj* function (73). The Time Course Inspector package was also used for calculating the mean and bootstrapped confidence intervals of replicate time series data. The image analysis pipeline, including all macros and R analysis scripts used for reporter normalizations and final replicate data processing are provided on the OSF project site (doi: 10.17605/OSF.IO/4F69N, components: *TIRF_analysis_pipeline*, *TIRF_datasets_Figs.1,2*).

### Live-cell epifluorescence microscopy of FOXO-based AKT kinase translocation reporter (KTR)

A detailed protocol of how HeLa cells were prepared for KTR measurements by live-cell widefield microscopy, including Matrigel coating of the dishes, cell seeding, and subsequent treatment has been made publicly available on protocols.io via the following doi: dx.doi.org/10.17504/protocols.io.261gedjkjv47/v1

Time-lapse epifluorescence images were obtained on a Nikon Ti2-E Inverted microscope fitted with a high-sensitivity CMOS Prime BSI (Photometric) sensor, with full temperature (37°C) and CO_2_ (5%) control throughout the acquisitions. A 10X 0.45 NA plan-apochromatic dry objective (Nikon) was used for illumination using the following set-up: LED-CFP/YFP/mCherry-3X-A Filter Cube; Triple Dichroic 459/526/596; Triple Emitter 475/543/702. Exposure times were 20 msec (for CLOVER) and 50 msec (for mCherry). Acquisition was performed in sequential mode, without binning, using an acquisition rate of 6 min.

Nuclear segmentation based on the NLS-mCherry fluorescence intensity was performed with Stardist in Fiji (74). Then, using custom-written Python scripts, the nuclear intensity *KTR*_*nuc*_was calculated as the average intensity of the KTR channel within a 5-by-5 pixel square around the centroid coordinates. Nuclear masks were then expanded by a width of 2 pixels, and the original mask was subtracted from the expanded one to generate the cytoplasmic ring mask. The cytoplasmic KTR intensity *KTR*_*cyto*_ was calculated as the average value of the brightest 50% pixels contained within this cytoplasmic ring mask. This was to avoid inclusion of background pixels in the calculations, in cases where cells were thin and elongated. The nuclear-to-cytoplasmic ratio *CN*_*R*_ was then computed as the ratio of cytoplasmic over total cellular intensities of the KTR sensor:

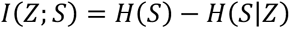

The *CN_R_* values were therefore bounded by 0 in the case of pure nuclear intensity (low pAKT levels), and 1 in the case of complete nuclear exclusion of the biosensor (high pAKT levels).

For trajectory generation, cells were first tracked using Trackmate (75) in Fiji, based on the centroids generated by the segmentation step. Tracks were filtered by length (only tracks persisting through the full time-course were conserved), and (x,y,t) coordinates from cellular tracks were matched with the corresponding *CN_R_* values using custom-written Python scripts. All subsequent data processing was performed in R for mean and bootstrapped confidence interval calculations (73), including visualization. All source data and scripts to reproduce the results are deposited on the OSF project site (doi: 10.17605/OSF.IO/4F69N, components *KTR_datasets_Figs.3,S3*, *KTR_datasets_2_Figs.3,S3, KTR_analysis_pipeline*).

### Scaffold-free spheroid generation and experimental processing

Scaffold-free spheroids were generated according to a new protocol developed in house for this study and made publicly available via the following doi: dx.doi.org/10.17504/protocols.io.3byl4bnrrvo5/v1. Spheroids were seeded at 1000-2000 cells/spheroid and used for experimentation 48h later. Prior to growth factor or inhibitor treatments, HeLa spheroids were serum-starved for 4h by an initial wash in 200 µl (96-well plate) or 3 ml (24-well plate) DMEM high-glucose (Thermo Fisher Scientific #41966-029) supplemented with another 2 mM of L-Glutamine. Human iPSC spheroids were washed and growth factor-depleted using DMEM/F12 (Thermo Fisher Scientific #21331-046) for 2 h prior to stimulation. In each case, the wash was removed and replaced with 100 µl (96-well plate) or 1 ml (24-well plate) of the same solution. Growth factor and inhibitor solutions were prepared as 3x working solutions and 50 µl (96-well plate) or 500 µl (24-well plate) of each added to the cells when required for a final dilution to 1x (1-100 nM for IGF1 or EGF; 500 nM for BYL719). Corresponding control solutions containing DMSO or non-DEPC-treated sterile water (Ambion #9937) were also applied. At the end of a time course, the spheroids were either processed for multiplexed mass cytometry (96-well plates) or RT-qPCR (24-well plates) as described below.

### Multiplexed mass cytometry (CyTOF) using 3D spheroids

A detailed step-by-step protocol for spheroid fixation, TOB*is* barcoding, enzyme-free single-cell dissociation and subsequent antibody staining for mass cytometry has been made publicly available via protocols.io: dx.doi.org/10.17504/protocols.io.4r3l22bz4l1y/v1. Final cell acquisition was performed either on a Helios or an XT mass cytometer, both developed by Fluidigm (now Standard Biotools). Raw mass cytometry data were normalized using bead standards (76) and debarcoded as per the computational algorithm developed by Zunder et al. (77). The Mahalanobis and separation cutoff were set to 10 and 0.1, respectively. Debarcoded .fcs files were imported into Cytobank (http://www.cytobank.org/) and gated with Gaussian parameters to remove debris, followed by gating on DNA (Ir-191/193), total S6, pRB^S807/S811^ and cCASP3^D175^ to separate cell populations according to cell state. For the iPSCs, there were no distinct populations of cCASP3^D175^-posite and -negative cells, which meant that this marker was not used for gating.

Gated populations of interest were exported as untransformed .txt files (excluding header with filename) and pre-processed using the CyGNAL package as described in Sufi et al. (28). The pre-processed .fcs files were imported into R as an object of the *SingleCellExperiment* class and *arcsinh* (inverse hyperbolic sine) transformed with cofactor = 5 using the flowCore (78) and CATALYST (79) packages in R. Individual marker histograms were generated using the ggridges R package. Exact gate settings, debarcoded and gated .fcs files as well as detailed scripts to reproduce all results are available on the OSF project site in dedicated subfolders (doi: 10.17605/OSF.IO/4F69N; component: *Mass_cytometry_CyTOF_HeLa_Figs.4,S4,S5,S6*). These subfolders also contain exact single-cell numbers for each experimental analysis in plots saved with the file suffix “_total_cell_count_plot.png”.

For EMD-PHATE analyses, the pre-processed .fcs files were imported into Python and processed using a custom-written script deposited on the OSF project site. The following markers were used for EMD-PHATE plot generation: “151Eu_pNDRG1 T346”, “167Er_pERK1_2_T202_Y204”, “173Yb_pS6_S240_S244”, “155Gd_pAKT S473”, “168Er_pSMAD2_3_S243_S245”.

### Reverse transcription-quantitative polymerase chain reaction (RT-qPCR)

Cellular RNA was extracted as described above for total mRNA Sequencing, and 250 ng used for complementary DNA (cDNA) synthesis with Thermo Fisher’s High-Capacity cDNA Reverse Transcription Kit (#4368814). Subsequent SYBR Green-based qPCRs were performed on 2.5 ng total cDNA.

A 5-fold cDNA dilution series was also prepared and used as standard curve for relative quantitation of gene expression. *TBP* was used as normalizer following confirmation that its gene expression was changing systematically as a function of the tested conditions, which was not the case for *ACTB* (tested as an additional housekeeping gene). Melt curve analyses and separate agarose gel electrophoresis confirmed amplification of the correctly-sized, single product by each primer pair. All primers had amplification efficiencies 95%-105%. Samples were loaded in duplicate in 384-well plates.

All qPCR data were acquired on a Quant Studio™ 6 Real-Time PCR System (Thermo Fisher Scientific). The thermocycling conditions (SYBR Green reactions) were as follows (ramp rate 1.6°C/s for all): 50°C for 2 min, 95°C for 10 min, 40 cycles at 95°C for 15 sec and 60°C for 1 min, followed by melt curve analysis (95°C for 15 sec, 60°C for 1 min, and 95°C for 15 min with ramp rate 0.075°C/sec). All relevant primer sequences are included in Table S4. Source data and scripts to reproduce the results have been deposted on the OSF project site (doi: 10.17605/OSF.IO/4F69N, component: *RT_qPCR_replicates_combined_Fig.5*).

### Evaluation of cellular morphology

HeLa cells were seeded at a density of 4,000 cells/well in black Perkin Elmer ViewPlate-96 dishes (TC-treated, #6005182) coated with Matrigel as per the protocol for TIRF and KTR imaging. After 24 h, 33 µl of 16 % methanol-free formaldehyde (Polysciences #18814-20) was added to 100 µl of culture medium in each well to fix the cells in 4% final formaldehyde concentration. Following 15 min of incubation at room temperature away from light, the fixative was removed and cells washed once with 100 µl DPBS, dispensed slowly and at a 45 degrees angle to prevent the cells from dislodging. Next, 75 µl of DPBS was added, followed by 19 µl of 5X fish skin gelatin blocking agent (Biotium #22010) diluted in PBS/T (PBS with 0.05 % Tween-20) with 0.5% Triton-X100 (5X concentration, diluted to 0.1 % once added to the DPBS). The cells were left to permeabilize and block for 10 min, after which the block/perm solution was removed and replaced with DPBS supplemented with Phalloidin iFlouor 555 (Abcam #ab176756) and HCS CellMask Blue (Thermo Fisher Scientific #H32720), both diluted 1:1000. Following 30 min incubation at room temperature away from light, the staining solution was removed and the cells washed twice with 100 µl DPBS. Another 100 µl of DPBS was added after the last wash, followed by epifluorescence acquisition on a Nikon Ti2 Eclipse microscope fitted with a 10X 0.45 NA plan-apochromatic dry objective (Nikon), used for imaging with the following set-up: LED-CFP/YFP/mCherry-3X-A Filter Cube; Triple Dichroic 459/526/596; Triple Emitter 475/543/702. Exposure times were 20 msec (for CellMaskBlue) and 60 msec (for Phalloidin and mCherry).

The CellMask blue images were converted to jpg for segmentation using Cellpose (v1) (80), and the resulting masks converted to regions of interests (ROIs) using the LabelsToROI plugin in Fiji/ImageJ. All edge ROIs were removed. The remaining ROIs were used for calculating the shape properties of the cell masks using the “Measure” function in Fiji/ImageJ.

Analysis of variance (ANOVA) models was fit in R to test for differences in solidity and area as a function of genotype. Of note, although the normality assumption was violated for these models, the impact of this is likely to be minimal given the large number of single cell observations and the assumptions of the central limit theorem. Tukey’s Honest Significant Differences method was used to test for statistically significant (adjusted p-value < 0.05) differences in shape properties as a function of genotype.

All raw images, segmentation masks, quantification scripts and a final montage of all composite images are deposited on the OSF project site (doi: 10.17605/OSF.IO/4F69N, component: *HeLa_cell_morphology_image_analysis_Fig.5*).

### Information theoretic analyses

For the estimation of mutual information and information capacity, the SLEMI (Statistical Learning-based Estimation of Mutual Information) R package was used (26). SLEMI uses a logistic regression model to learn the discrete probability *P*(*S*|*Z*) of the signal (S) given the response (Z) and subsequently estimates the mutual information, *I*(*Z*; *S*), from the following formula:

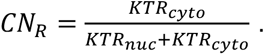

Here, *H*(*S*) = − ∑*_s_ P*(*s*)log_2_(*P*(*s*)) is the entropy of the signal calculated based on the input signal distribution *P*(*S*); and *H*(*S*|*Z*) = *E*[− ∑*_s_ P*(*s*|*Z*)log_2_(*P*(*s*|*Z*))] is the conditional entropy of the signal given the response. A uniform signal distribution was used in all cases for mutual information estimation, while information capacity is estimated by maximizing the mutual information over possible signal distributions. This approach does not rely on any form of data binning and is therefore particularly well-suited for the case of high-dimensional outputs such as the time course measurements in our live cell experiments.

The specific scripts used to calculate mutual information are deposited on the OSF project site (doi: 10.17605/OSF.IO/4F69N, components: *TIRF_datasets_Figs.1,2*, *KTR_datasets_2_Figs.3,S3*).

### Statistics and reproducibility

Bespoke data and statistical analyses are detailed in the relevant methods sections. Note that the information theoretic analyses detailed above also provide the appropriate statistical description of the trajectory datasets as justified by Bayesian decision theory (81). The mass cytometry datasets are shown as individual distributions, with probability-based thresholding as opposed to use of metrics such as standard deviation and variance given the non-normal distribution of the data. In general, rather than applying conventional statistical tests that would be violated by the structure of our data, we chose to focus on orthogonal validation in independent model systems.

Source data and annotated scripts to reproduce all results are included on the OSF project site (doi: 10.17605/OSF.IO/4F69N).

## Acknowledgements

We would like to thank the following colleagues and collaborators for technical and scientific advice, including protocol sharing and helpful discussions: Dr. Gerry Hammond (University of Pittsburgh), Dr. James Burchfield (University of Sydney), Dr. Alison Kearney (University of Sydney), Dr. James Opzoomer (UCL Cancer Institute), Dr. Elitza Deltcheva (UCL Cancer Institute), Dr. Benoit Bilanges (UCL Cancer Institute), Prof. Alex Toker (Beth Israel Deaconess Medical Centre), Prof. Andre Levchenko (Yale University) and Prof. Robert Semple (University of Edinburgh). We are indebted to the following core facilities and their staff for technical assistance throughout this project: UCL BLIC (Dr. Lucia Conde), UCL Cancer Institute Flow Cytometry Facility, UCL Cancer Institute Imaging Facility, Dundee Imaging Facility, and The Francis Crick Institute Flow Cytometry Facility. We are also grateful to members of the Payne Lab (UCL Cancer Institute) for their assistance with MiSeq library processing.

## Author contributions

R.R.M. conceived and supervised the project, acquired funding, conducted experiments, analyzed data, developed analytical pipelines, prepared figures and wrote the manuscript. A.L.M. conducted KTR experiments and developed the associated analytical pipeline. O.M. conducted KTR experiments, analyzed data, developed an open-source version of the KTR analysis pipeline and provided routine technical assistance. M.V. performed all information theoretic analyses. S.Y. conducted TIRF and RT-qPCR experiments, analyzed data and provided routine technical assistance. D.M., J.S., S.J.Z., J.G. and L.D. provided technical assistance. X.Q. wrote the EMD-PHATE analysis code. B.V. helped with project supervision, acquired funding and provided extensive feedback on the first manuscript draft. C.T. helped with project supervision and acquired funding. E.H. acquired funding. V.K. helped with project supervision. All authors reviewed the final manuscript.

## KEY RESOURCES TABLES

**Table 1:** Primary Antibodies for Western blotting.

| Antibody target | Clone | Mol. Weight (kDa) | Lot # | Vendor | Cat. # | Dilution |
| --- | --- | --- | --- | --- | --- | --- |
| IGF1Rβ (C-20) | C-20 | 95 | J1907 | Santa Cruz | sc-713 | 1:200 |
| INSRβ (C-20) | C-20 | 95 | L1812 | Santa Cruz | sc-711 | 1:200 |
| EGFR | Polyclonal | 175 | 17 | CST | 2232 | 1:1000 |
| p110α | C73F8 | 110 | 11 | CST | 4249 | 1:1000 |
| p110β | C33D4 | 110 | 8 | CST | 3011 | 1:1000 |
| pAKT T308 | 244F9 | 60 | 17 | CST | 4056 | 1:1000 |
| pAKT Ser473 | Polyclonal | 60 | 14 | CST | 9271 | 1:1000 |
| Total AKT | Polyclonal | 60 | 28 | CST | 9272 | 1:1000 |
| pERK1/2 (T202; Y204) | Polyclonal | 44/42 | 17 | CST | 4370 | 1:1000 |
| pS6 (S240/S244) | Polyclonal | 32 | 18 | CST | 2215 | 1:1000 |
| Total S6 | 54D2 | 32 | 13 | CST | 2317 | 1:1000 |
| Vinculin |  | 124 | NA | Sigma Aldrich | V9131 | 1:5000 |

**Table 2:** Secondary Antibodies for Western blotting.

| Antibody | Vendor | Lot # | Cat. # | Dilution |
| --- | --- | --- | --- | --- |
| anti-rabbit HRP-conjugated | Amersham | 17203153 | NA934V | 1:5000 |
| anti-mouse HRP-conjugated | Amersham | 17193521 | NXA931V | 1:5000 |
| Goat anti-rabbit IgG HRP-linked antibody | CST | 30 | 7074S | 1:10000 |
| Goat anti-mouse IgG HRP-linked antibody | CST | 36 | 7076S | 1:10000 |

**Table 3:** Mass cytometry antibodies.

| Antibody target | Clone | Metal | Lot # | Vendor | Cat. # | Amount (1 rxn) |
| --- | --- | --- | --- | --- | --- | --- |
| Total S6 | 54D2 | 141-Pr | 16 | CST | 2317 | 0.32 µg |
| Cleaved Caspase 3 (D175) | D3E9 | 142-Nd | 4 | CST | 9579 | 0.59 µg |
| pRB (S807/811) | J112-906 | 150-Nd | 2005420 | Fluidigm | 3150013 | 0.7 µg |
| pNDRG1 (T346) | D98G11 | 151-Eu | 4 | CST | 5482 | 0.56 µg |
| pAKT (S473) | M89-16 | 155-Gd | 20 | BD Biosciences | 560397 | 4 µg |
| pERK1/2 (T202/Y204) | 20A | 167-Er | 1263406 | BD Biosciences | 612359 | 0.09 µg |
| pSMAD2/3 (S263/S265; S423/S425) | D27F4 | 168-Er | 2 | CST | 8828 | 1.72 µg |
| pS6 (S240/S244) | D68F8 | 173-Yb | 7 | CST | 5364 | 0.46 µg |

**Table 4:**
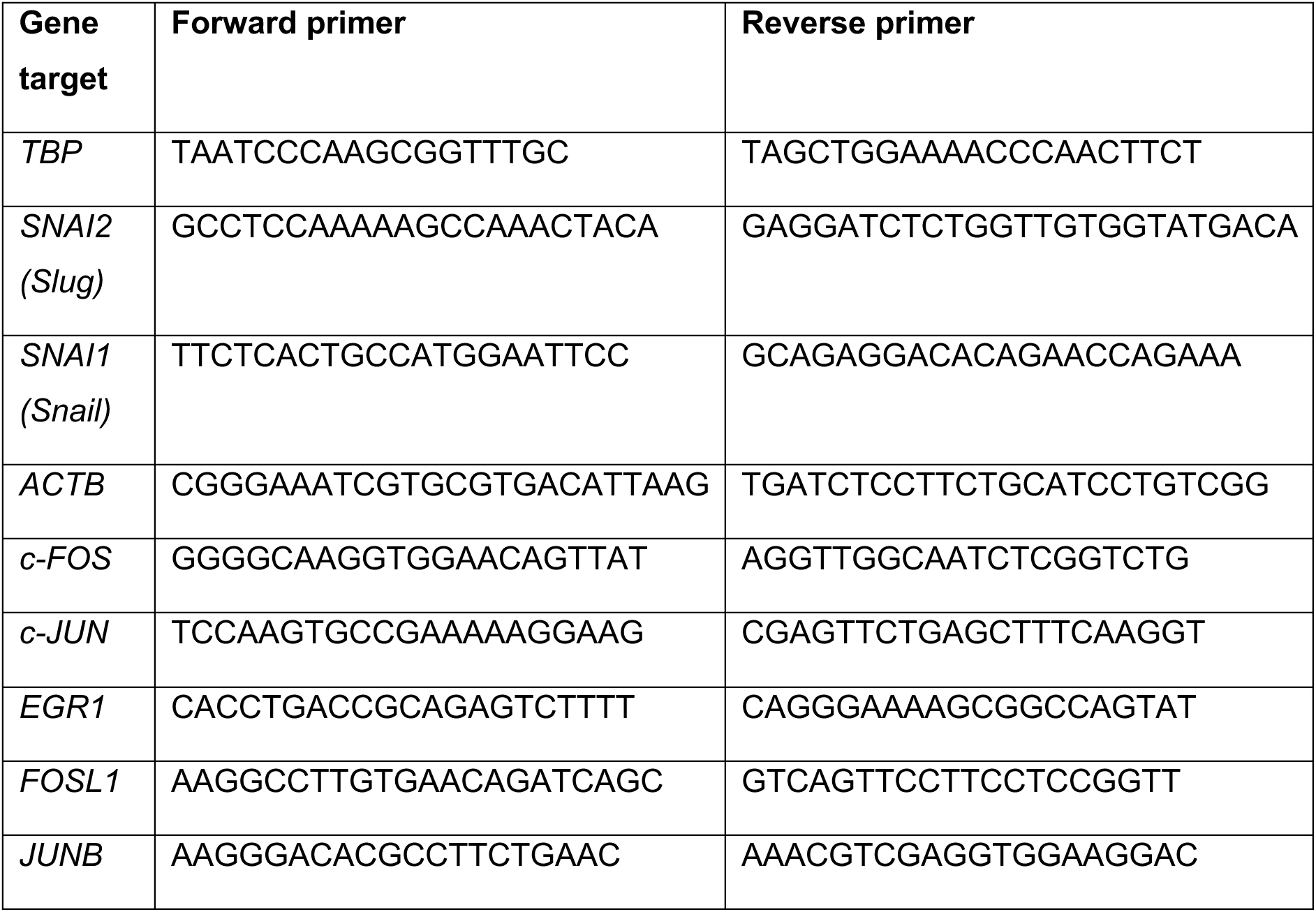
RT-qPCR Primers.

**Table 5:** Cell Lines.

| Cell line | Source |
| --- | --- |
| HeLa cervical cancer cells (*PIK3CA* WT, 1xH1047R, 2xH1047R) | Engineered as part of this work, from parental ATCC HeLa line (#CCL-2) |
| Immortalised mouse embryonic fibroblasts (*PIK3CA* WT or Cre-del *PIK3CA*-null) | In house (see Ref. (82)) |
| CRISPR/Cas9-engineered A549 lung adenocarcinoma cells (*PIK3CA*-WT or *PIK3CA*-null) | In house (see Ref. (15)) |
| CRISPR/Cas9-engineered WTC11 human induced pluripotent stem cells (*PIK3CA* WT/WT, WT/H1047R, H1047R/H1047R) | A gift from Professor Robert Semple, University of Edinburgh (see Ref. (24)) |

**Table 7:**
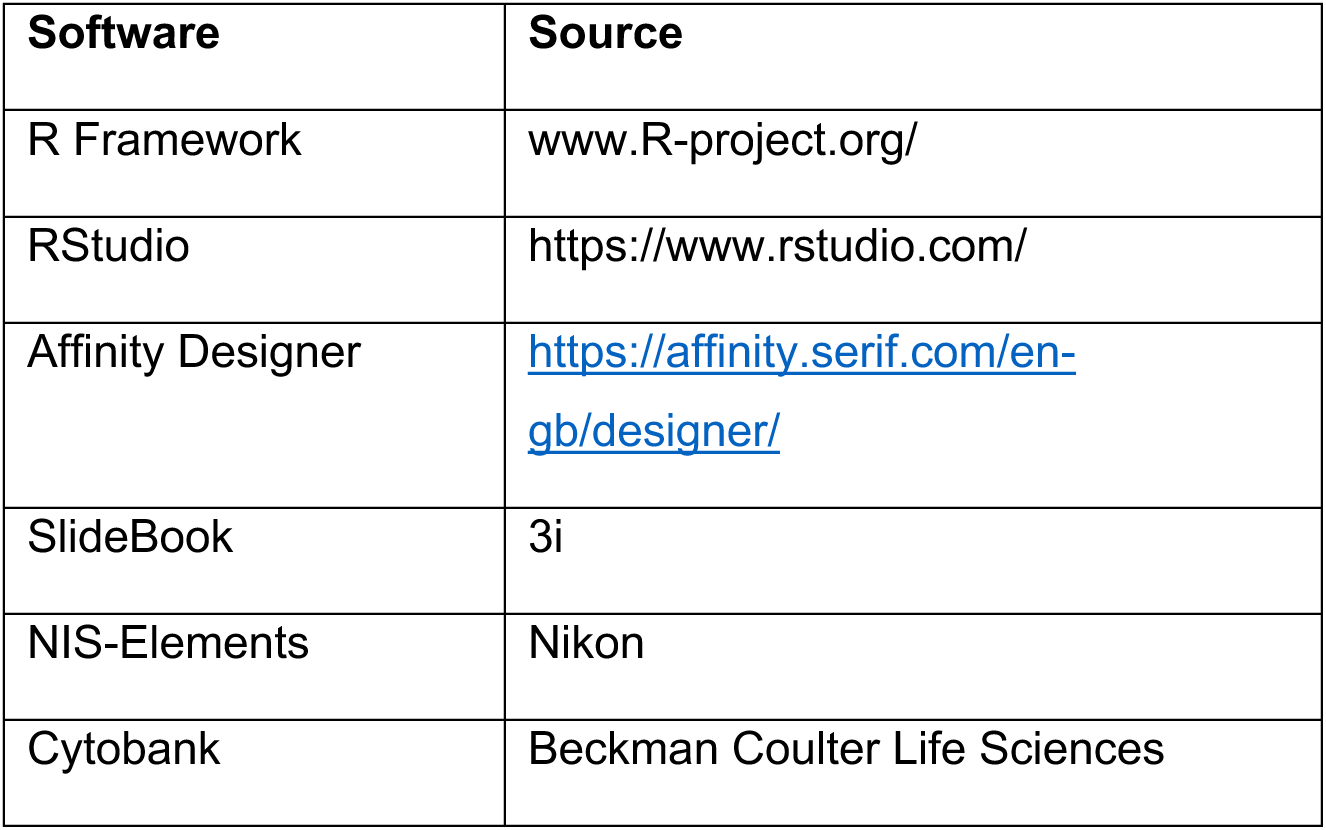
Software & Algorithms.

| Software | Source |
| --- | --- |
| R Framework | www.R-project.org/ |
| RStudio | https://www.rstudio.com/ |
| Affinity Designer | https://affinity.serif.com/en-gb/designer/ |
| SlideBook | 3i |
| NIS-Elements | Nikon |
| Cytobank | Beckman Coulter Life Sciences |

## SUPPLEMENTARY MATERIAL

Additional source data and all annotated analysis workflows are available via a bespoke OSF project website (doi: 10.17605/OSF.IO/4F69N).

Detailed protocols for spheroid, TIRF, KTR and mass cytometry experimental set-ups are provided on protocols.io via the following doi links:

□ Spheroid set-up: dx.doi.org/10.17504/protocols.io.3byl4bnrrvo5/v1
□ Preparation of cells for live-cell imaging of phosphoinositide reporters by total internal reflection fluorescence (TIRF) microscopy: dx.doi.org/10.17504/protocols.io.kxygx37jkg8j/v1
□ Preparation of cells for live-cell imaging of a FOXO-based AKT kinase translocation reporter by widefield microscopy: dx.doi.org/10.17504/protocols.io.261gedjkjv47/v1
□ Processing of fixed spheroids for TOBis barcoding, enzyme-free dissociation and antibody staining for CyTOF: dx.doi.org/10.17504/protocols.io.4r3l22bz4l1y/v1

**Figure S1.**
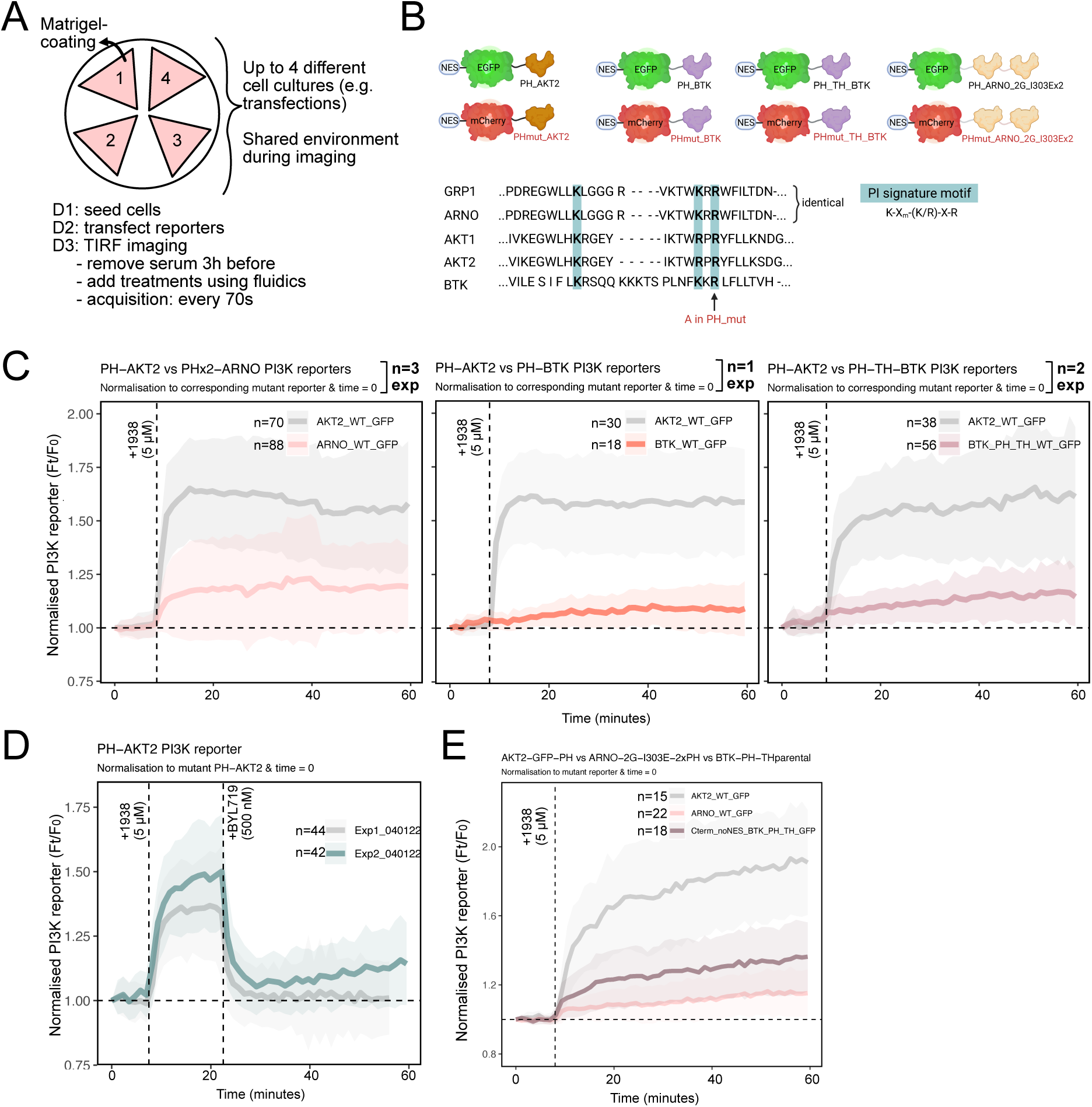
Systematic benchmarking of pleckstrin homology (PH) domain-based class I PI3K biosensors. **(A)** Schematic of the optimized live-cell imaging set-up to ensure that multiple comparisons could be performed in the same microenvironment, aided by fluidics for minimal physical perturbation during compound additions. To bring down the baseline of PI3K signaling, serum was removed from the cells 3 h prior to imaging start. D1, D2, D3 refer to day 1, day 2 and day 3 of the experimental workflow. **(B)** Schematic of the different wild-type and mutant PH domains used for benchmarking, all cloned into the same plasmid backbone for consistent comparisons. The PH domain of GRP1 has often been used for live-cell detection of PIP_3_ in the literature. Its sequence in the region responsible for PIP_3_ is identical to the ARF GEF ARNO. We therefore chose to include the latter in our comparisons given the recently reported a tandem-dimer, modified version of this PH domain as an improved biosensor for PIP_3_ (18).The shown alignments cover the conserved *β1* strand, variable loop 1, and *β2* strand of the PH domain fold. Of the four PH domains, only PH-AKT2 is capable of binding both PIP_3_ and PI(3,4)P_2_. The remaining PH domains only bind PIP_3_. The Alanine (A) mutation in the phosphoinositide (PI) signature motif renders the mCherry-tagged mutant PH domain versions unable to bind phosphoinositides. **(C)** Quantification of total internal reflection fluorescence (TIRF) microscopy experiments comparing the response rate and dynamic range of individual PH domain-based PI3K reporters in response to pharmacological PI3Kα activation in HeLa cells. To correct for non-specific increases in biosensor signal at the plasma membrane, the intensity of each GFP-tagged wild-type PH domain was normalized to that of its mCherry-tagged mutant version. The traces represent mean fold-change relative to baseline (the median signal of the first four time points), with shaded areas representing +/- 1 standard deviation (SD). Experimental replicates and single-cell numbers are indicated. Two different constellations were tested for BTK-derived PH domain: with and without the adjacent Tec homology (TH) domain. Only one experiment was performed with PH-BTK without TH as most of the cells failed to tolerate its expression. **(D)** TIRF-M of the PH-AKT2-derived biosensor in HeLa cells stimulated with 5 µM 1938, followed by pharmacological PI3Kα inhibition with 500 nM BYL719. Two independent experiments are superimposed to illustrate the expected inter-experimental variability. All plots represent mean normalized reporter signal relative to time 0, with shading corresponding to standard deviation. **(E)** The performance of the PH-TH version of BTK could be improved when switching from N-terminal to C-terminal fluorescent protein fusion, with simultaneous removal of the nuclear export sequence, as in the original plasmid DNA used for subcloning of this reporter.

**Fig. S2.**
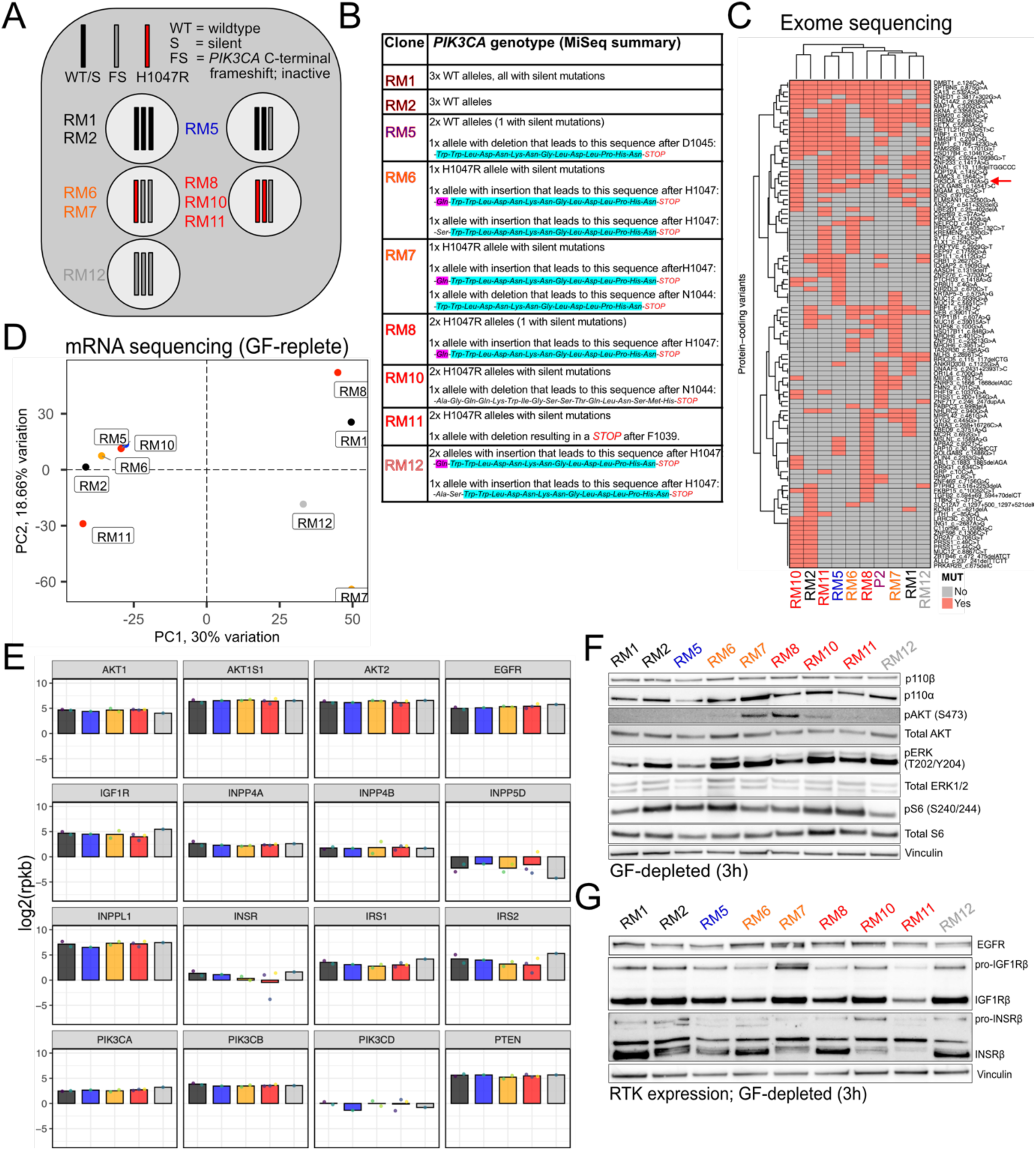
CRISPR/Cas9 *PIK3CA* exon21 engineering and quality control assays of HeLa clones. **(A)** Summary of the final set of clones that were banked and validated following engineering. As HeLa cells are nominally triploid, the edited clones either have three wildtype *PIK3CA* alleles or either one of the following: 0-2 wildtype *PIK3CA* alleles, 0-2 *PIK3CA^H1047R^* alleles and/or 0-3 C-terminal frameshift alleles. **(B)** The C-terminal frameshift truncation results in recoding of the last c. 20-30 amino acids of the p110α protein (common reading frames are highlighted in purple and turquoise). This abolishes the critical p110α WIF motif required for membrane binding and catalytic function (38), effectively creating a loss-of-function knock-in that does not carry the risk of a complete knock-out in terms of altering the stoichiometry of regulatory and catalytic p110 subunits. The exact allelic sequence was obtained following targeted next generation sequencing of the edited *PIK3CA* exon21 region. **(C)** Clustering of individual HeLa cultures based on protein-coding gene variants found in two or more CRISPR/Cas9 engineered HeLa clones relative to the parental culture prior to gene editing. Gene variants were captured using whole exome sequencing, which included a control HeLa culture passaged alongside the CRISPR/Cas9-edited clones without any editing or subcloning. A red arrow is used to indicate the *PIK3CA^H1047R^* edit which is the only protein-coding variant common to all *PIK3CA^H1047R^* mutant cell lines. **(D)** Principal component analysis (PCA) of CRISPR/Cas9-engineered HeLa clones based on total mRNA sequencing data obtained in baseline culture condition following fresh medium replenishment 3 h prior to sample collection. The observed clustering is similar to that observed with the exome sequencing data in (C), without any systematic differences driven by the presence of the *PIK3CA^H1047R^* variant. **(E)** Barplots comparing the expression levels in log2(reads per kilobase) for selected PI3K pathway-relevant genes and receptor tyrosine kinases. **(F)** Western blots for PI3K signaling components and pERK1/2 following 3 h of serum or growth factor (GF) removal, using all CRISPR/Cas9-engineered HeLa clones. **(G)** As in F but focusing on the expression levels of relevant receptor tyrosine kinases. EGFR, epidermal growth factor receptor; IGF1R, insulin-like growth factor 1 receptor; INSRβ, insulin receptor β chain.

**Figure S3.**
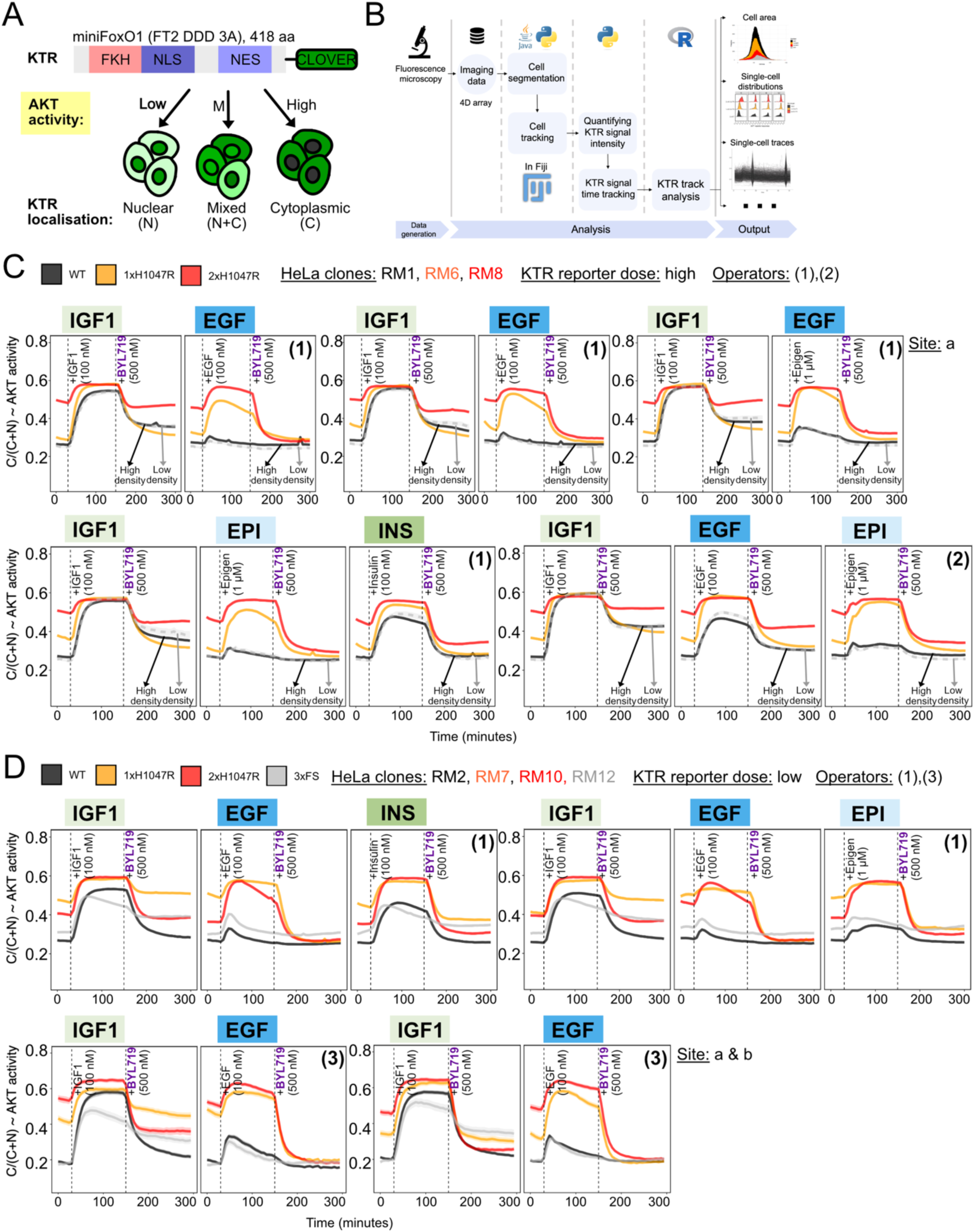
FOXO-based AKT kinase translocation reporter (KTR) set-up and full set of experimental outputs. **(A)** Schematic of the reporter and its mechanism of action. Note that the reporter was delivered into cells using transposon-based technology for stable expression. **(B)** Overview of the computational image and KTR data analysis pipeline which has been deposited on the accompanying OSF project site (doi: 10.17605/OSF.IO/4F69N). The data in **(C)** and **(D)** are from all independent experiments performed across different genotypes, HeLa clones, cell densities, KTR reporter doses, operators and experimental sites for a robust evaluation of reproducibility. For each time point, the traces correspond to the mean proportion of cytoplasmic KTR signal, with shaded areas representing bootstrapped 95% confidence intervals of the mean (note that these may be too small to be seen on the figure). Note that we observed operator-dependent differences in EGF-induced signaling dynamics in wild-type cells, yet the overall pattern relative to IGF1, including the blurring of the response in mutant cells, remained consistent. This technical variability in EGF responses in wild-type cells is likely due to their sensitivity to the pressure/rate of delivery of the stimulus through the manual fluidics system (see doi: dx.doi.org/10.17504/protocols.io.261gedjkjv47/v1).

**Fig. S4.**
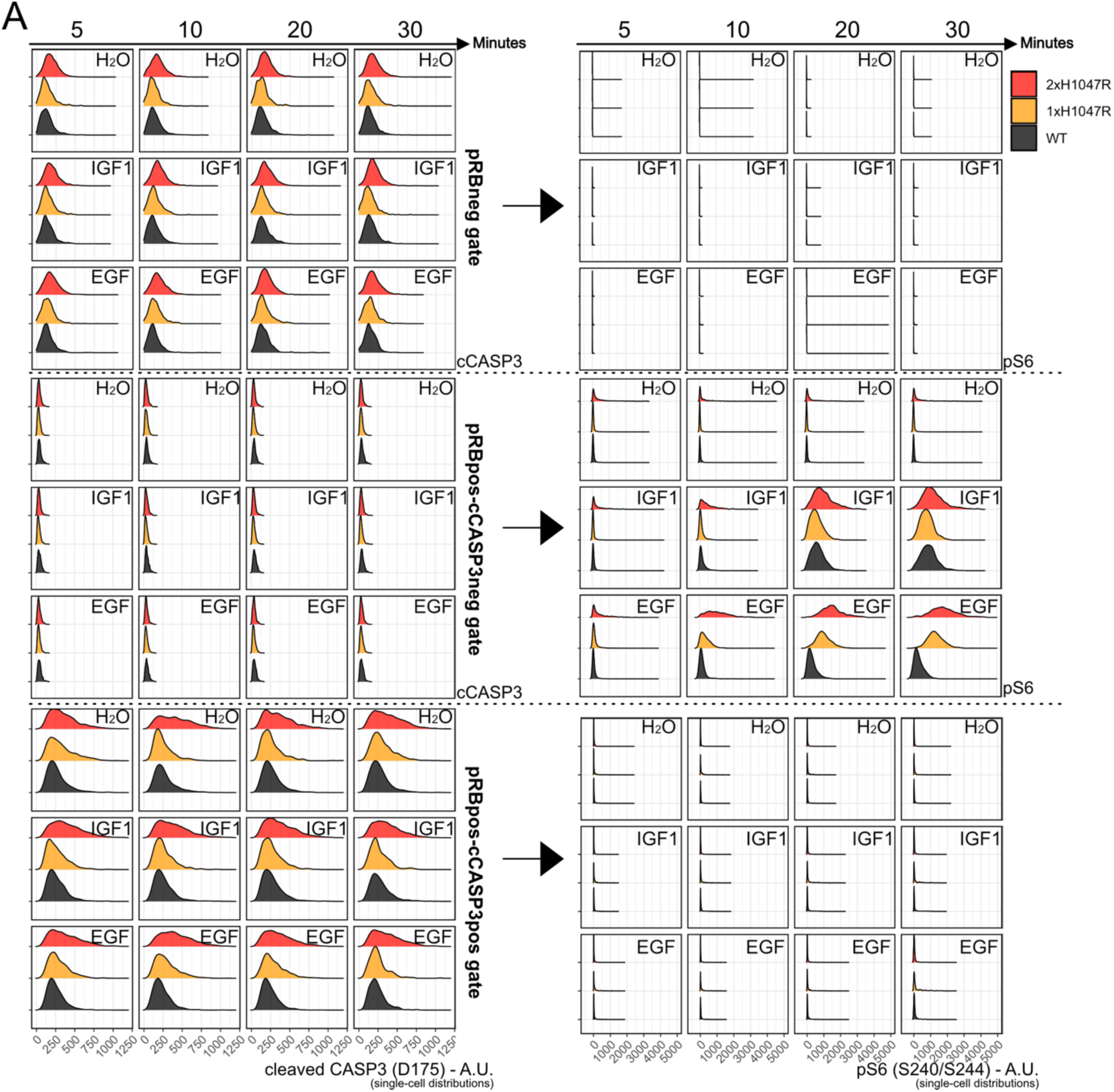
Example data demonstrating that growth factor-induced signaling responses are only observed in pRB^S807/S811^-positive (cycling) and cleaved Caspase^D175^-negative (non-apoptotic) HeLa spheroid cells. The plots on the left-hand side show the single-cell cCASP3^D175^ signal in the different pRB gates as indicated. The plots on the right show the corresponding pS6^S240/S244^ signal in each gate. The overall experimental setup is as shown in Fig. 4.

**Figure S5.**
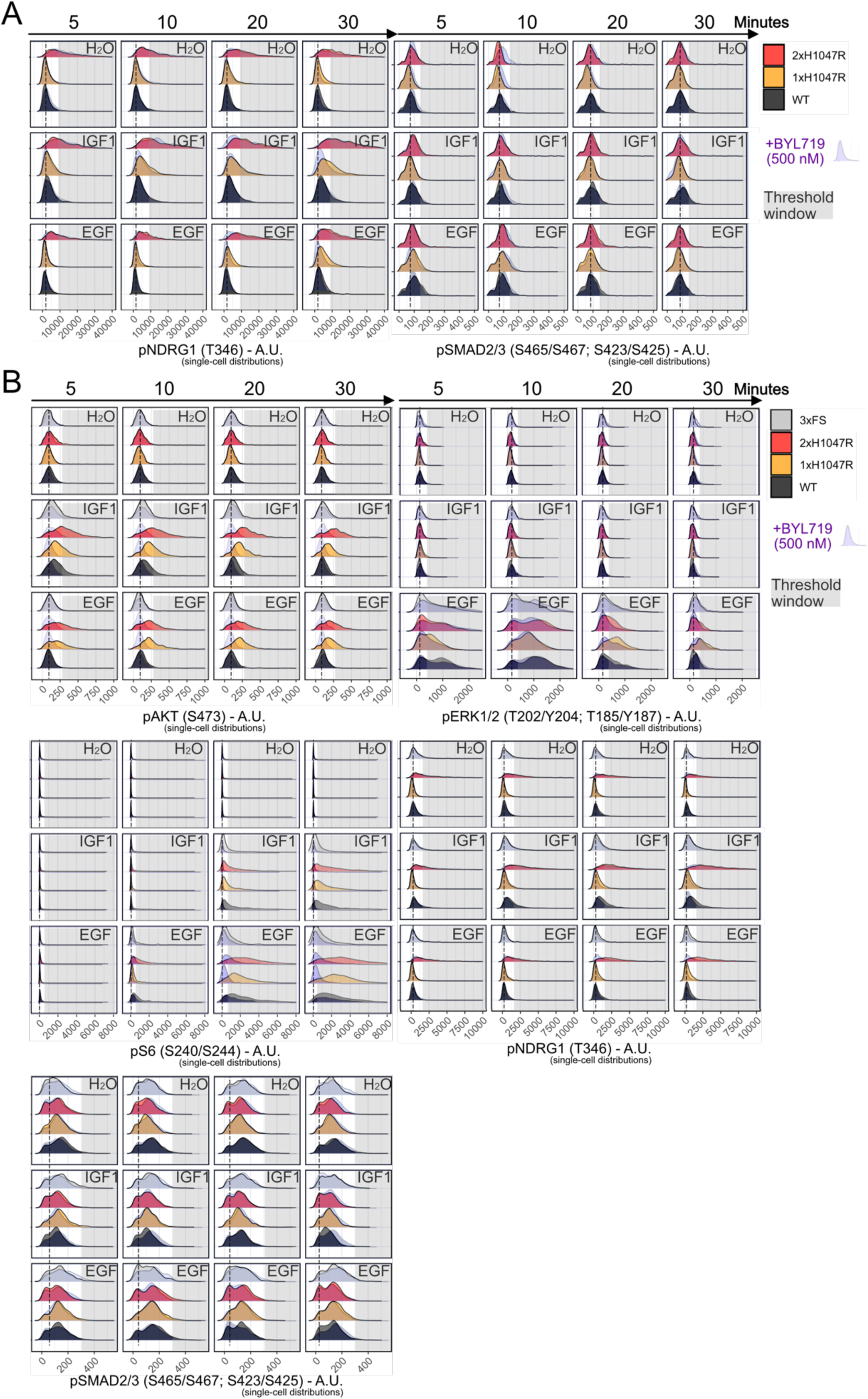
Additional CyTOF data and independent experimental replicate with independent clones. **(A)** The pNDRG1 and pSMAD2/3 signaling responses captured as part of the dataset shown in Fig. 4. The phosphorylation of NDRG1 on T346 is a known marker of mTORC2 activation (84). The phosphorylation of SMAD2/3 (S465/S467; S423/S425) is a marker for activated TGFβ signaling which is associated with *PIK3CA^H1047R^*phenotypes in human iPSCs (25). **(B)** CyTOF data from a repeat of the experiment in Fig. 4, using independent CRISPR/Cas9-engineered, 3D-cultured HeLa clones, including the *PIK3CA* loss-of-function 3xFS clone as an additional control. The spheroids were serum-starved for 4 h prior to the indicated perturbations. The shown signaling data are from cycling, non-apoptotic cells. The stippled line indicates the position of the peak in wild-type spheroids treated with vehicle (H_2_O). The grey shading highlights the response region that is not accessible to *PIK3CA* wild-type cells in the absence of stimulation.

**Figure S6.**
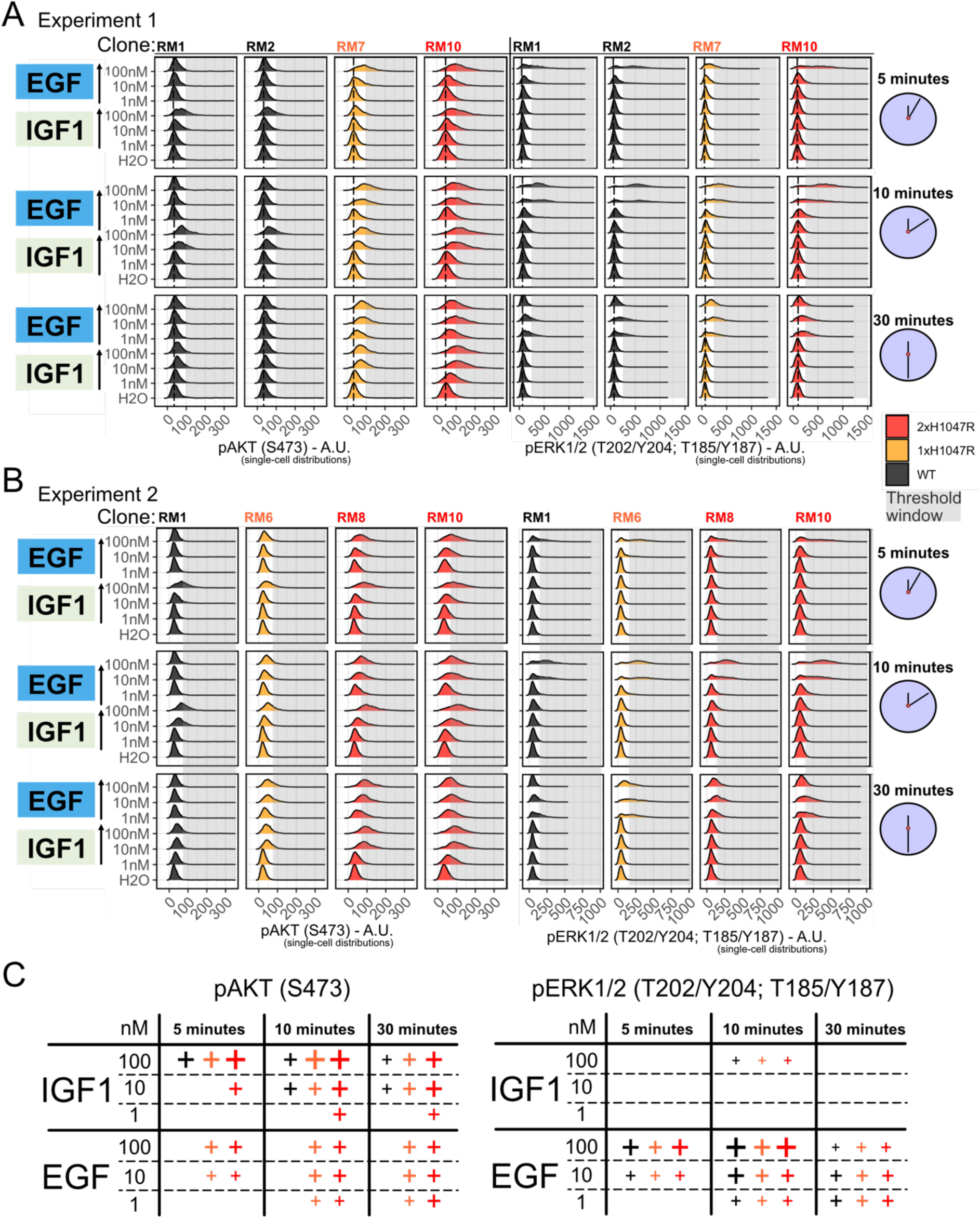
Dose- and time-dependent IGF1 and EGF single-cell signaling responses in HeLa spheroid cells with wild-type or *PIK3CA^H1047R^* (1-2 copies) expression. The plots in **(A)** and **(B)** are from two independent CyTOF datasets using independent CRISPR/Cas9-engineered, 3D-cultured HeLa clones stimulated with 1, 10 or 100 nM of IGF1 or EGF as a function of time. The spheroids were serum-starved for 4 h prior to the indicated perturbations. The shown signaling data are from cycling, non-apoptotic cells. The stippled line indicates the position of the peak in wild-type spheroids treated with vehicle (H_2_O). The grey shading highlights the response region that is not accessible to *PIK3CA* wild-type cells in the absence of stimulation. **(C)** Graphical summary of the key observations in the datasets in (A) and (B). A positive response is indicated with (+), the size of which indicates the response magnitude.

**Figure S7.**
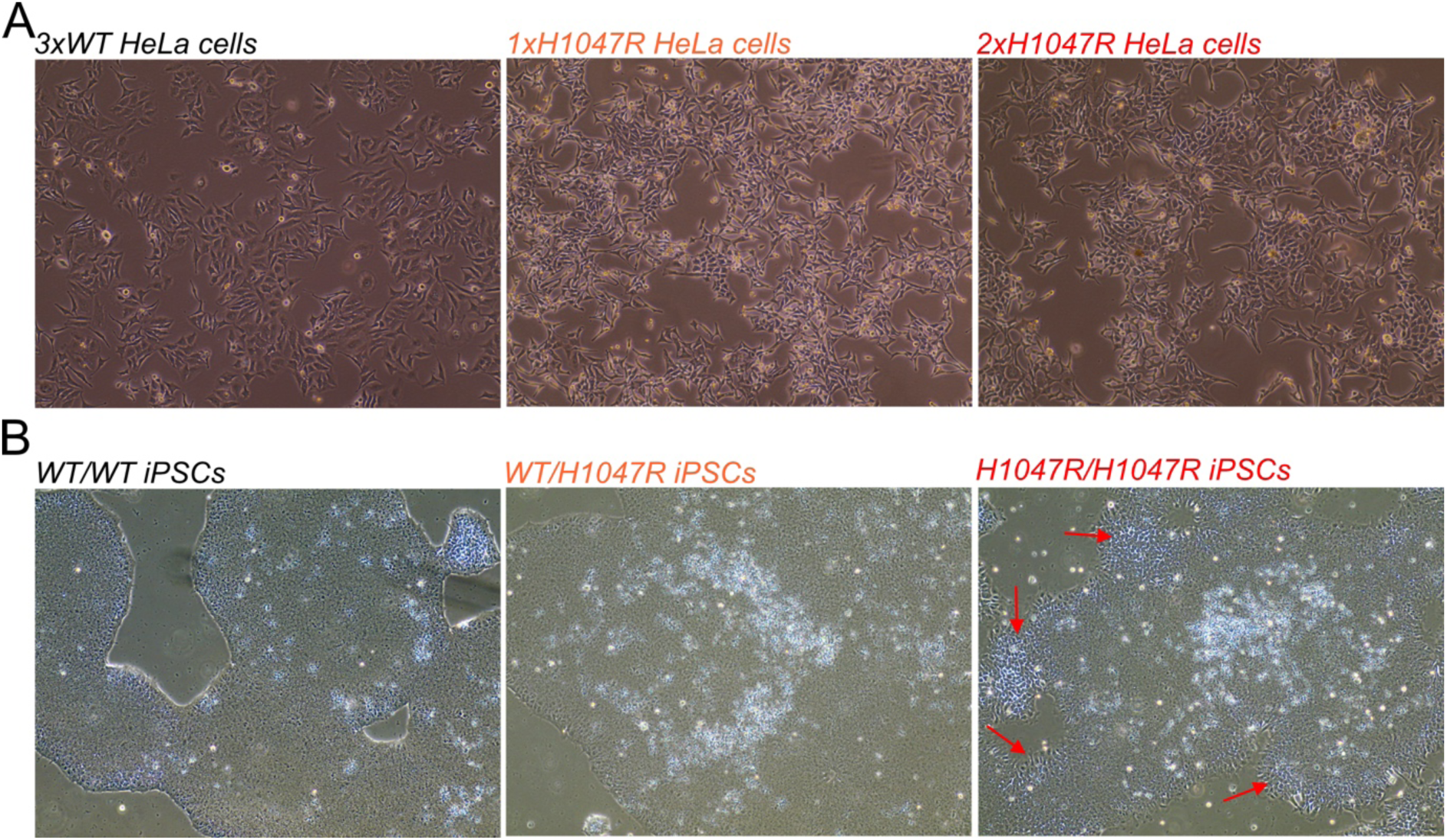
Light microscopy images of HeLa (A) and iPSC (B) maintenance cultures with different *PIK3CA* genotypes. The cultures in A correspond to independent CRISPR/Cas9-engineered HeLa clones to those used for staining in Fig. 5B. The red arrows in (B) point to example regions with disorganized colony growth and mesenchymal-like morphology changes in homozygous *PIK3CA^H1047R^* iPSCs.

## References

1. Isakoff SJ, Engelman JA, Irie HY, Luo J, Brachmann SM, Pearline RV, et al. Breast cancer-associated PIK3CA mutations are oncogenic in mammary epithelial cells. Cancer Research. 2005;65(23):10992–1000.

2. Samuels Y, Diaz LA, Schmidt-Kittler O, Cummins JM, DeLong L, Cheong I, et al. Mutant PIK3CA promotes cell growth and invasion of human cancer cells. Cancer Cell. 2005;7(6):561–73.

3. Kang S, Bader AG, Vogt PK. Phosphatidylinositol 3-kinase mutations identified in human cancer are oncogenic. Proceedings of the National Academy of Sciences. 2005;102(3):802–7.

4. Vanhaesebroeck B, Perry MWD, Brown JR, André F, Okkenhaug K. PI3K inhibitors are finally coming of age. Nature Reviews Drug Discovery. 2021 Oct 14;20(10):741–69.

5. Madsen RR, Vanhaesebroeck B, Semple RK. Cancer-Associated PIK3CA Mutations in Overgrowth Disorders. Trends in Molecular Medicine. 2018 Oct;24(10):856–70.

6. Canaud G, Gutierrez JCL, Irvine AD, Vabres P, Hansford JR, Ankrah N, et al. Alpelisib for Treatment of Patients With PIK3CA-Related Overgrowth Spectrum (PROS). Genetics in Medicine. 2023 Aug;100969.

7. Madsen RR, Vanhaesebroeck B. Cracking the context-specific PI3K signaling code. Science Signaling. 2020 Jan 7;13(613):eaay2940.

8. Madsen RR, Toker A. PI3K signaling through a biochemical systems lens. Journal of Biological Chemistry. 2023 Oct;299(10):105224.

9. Bugaj LJ, Sabnis AJ, Mitchell A, Garbarino JE, Toettcher JE, Bivona TG, et al. Cancer mutations and targeted drugs can disrupt dynamic signal encoding by the Ras-Erk pathway. Science. 2018 Aug 31;361(6405):eaao3048.

10. Levchenko A, Nemenman I. Cellular noise and information transmission. Current Opinion in Biotechnology. 2014 Aug;28:156–64.

11. Cheong R, Rhee A, Wang CJ, Nemenman I, Levchenko A. Information transduction capacity of noisy biochemical signaling networks. Science. 2011 Oct 21;334(6054):354–8.

12. Selimkhanov J, Taylor B, Yao J, Pilko A, Albeck J, Hoffmann A, et al. Accurate information transmission through dynamic biochemical signaling networks. Science. 2014 Dec 12;346(6215):1370–3.

13. Chen JY, Lin JR, Cimprich KA, Meyer T. A Two-Dimensional ERK-AKT Signaling Code for an NGF-Triggered Cell-Fate Decision. Molecular Cell. 2012;45(2):196–209.

14. Levchenko A. Genetic diseases: How the noise fits in. Current Biology. 2023 Mar 27;33(6):R228–30.

15. Gong GQ, Bilanges B, Allsop B, Masson GR, Roberton V, Askwith T, et al. A small-molecule PI3Kα activator for cardioprotection and neuroregeneration. Nature. 2023 May 24;1–10.

16. Vihinen M, Nilsson L, Smith CIE. Tec homology (TH) adjacent to the PH domain. FEBS Letters. 1994;350(2–3):263–5.

17. Chung JK, Nocka LM, Decker A, Wang Q, Kadlecek TA, Weiss A, et al. Switch-like activation of Bruton’s tyrosine kinase by membrane-mediated dimerization. Proceedings of the National Academy of Sciences. 2019;201819309.

18. Goulden BD, Pacheco J, Dull A, Zewe JP, Deiters A, Hammond GRV. A high-avidity biosensor reveals plasma membrane PI(3,4)P2 is predominantly a class I PI3K signaling product. Journal of Cell Biology. 2019;218(3):1066–79.

19. Ebner M, Lučić I, Leonard TA, Yudushkin I. PI(3,4,5)P3 Engagement Restricts Akt Activity to Cellular Membranes. Molecular Cell. 2017;65(3):416–431.e6.

20. Cronin TC, DiNitto JP, Czech MP, Lambright DG. Structural determinants of phosphoinositide selectivity in splice variants of Grp1 family PH domains. EMBO Journal. 2004;23(19):3711–20.

21. Bekker-Jensen DB, Kelstrup CD, Batth TS, Larsen SC, Haldrup C, Bramsen JB, et al. An Optimized Shotgun Strategy for the Rapid Generation of Comprehensive Human Proteomes. Cell Systems. 2017;4(6):587–599.e4.

22. Saito Y, Koya J, Araki M, Kogure Y, Shingaki S, Tabata M, et al. Landscape and function of multiple mutations within individual oncogenes. Nature. 2020 Jun 8;582(7810):95–9.

23. Sivakumar S, Jin DX, Rathod R, Ross J, Cantley LC, Scaltriti M, et al. Genetic heterogeneity and tissue-specific patterns of tumors with multiple *PIK3CA* mutations. Clinical Cancer Research. 2023 Jan 3;CCR-22-2270.

24. Madsen RR, Knox RG, Pearce W, Lopez S, Mahler-Araujo B, McGranahan N, et al. Oncogenic PIK3CA promotes cellular stemness in an allele dose-dependent manner. Proceedings of the National Academy of Sciences. 2019 Apr 23;116(17):8380–9.

25. Madsen RR, Longden J, Knox RG, Robin X, Völlmy F, Macleod KG, et al. NODAL/TGFβ signalling mediates the self-sustained stemness induced by PIK3CAH1047R homozygosity in pluripotent stem cells. Disease Models & Mechanisms. 2021 Mar 1;14(3):dmm.048298.

26. Jetka T, Nienałtowski K, Winarski T, Błoński S, Komorowski M. Information-theoretic analysis of multivariate single-cell signaling responses. Rao CV, editor. PLoS Comput Biol. 2019 Jul 12;15(7):e1007132.

27. Gross SM, Dane MA, Bucher E, Heiser LM. Individual Cells Can Resolve Variations in Stimulus Intensity along the IGF-PI3K-AKT Signaling Axis. Cell Systems. 2019 Dec;9(6):580–588.e4.

28. Sufi J, Qin X, Rodriguez FC, Bu YJ, Vlckova P, Zapatero MR, et al. Multiplexed single-cell analysis of organoid signaling networks. Nature Protocols. 2021 Sep 8;16(10):4897– 918.

29. Kramer BA, Sarabia del Castillo J, Pelkmans L. Multimodal perception links cellular state to decision-making in single cells. Science. 2022 Aug 5;377(6606):642–8.

30. Moon KR, van Dijk D, Wang Z, Gigante S, Burkhardt DB, Chen WS, et al. Visualizing structure and transitions in high-dimensional biological data. Nature Biotechnology. 2019;37(12):1482–92.

31. Avraham R, Yarden Y. Feedback regulation of EGFR signalling: decision making by early and delayed loops. Nat Rev Mol Cell Biol. 2011 Feb;12(2):104–17.

32. Ram A, Murphy D, DeCuzzi N, Patankar M, Hu J, Pargett M, et al. A guide to ERK dynamics, part 2: downstream decoding. Biochemical Journal. 2023 Dec 1;480(23):1909–28.

33. Cook DP, Vanderhyden BC. Context specificity of the EMT transcriptional response. Nature communications. 2020;11(1):2142.

34. Devaraj V, Bose B. Morphological State Transition Dynamics in EGF-Induced Epithelial to Mesenchymal Transition. Journal of Clinical Medicine. 2019 Jul;8(7):911.

35. Gustin JP, Karakas B, Weiss MB, Abukhdeir AM, Lauring J, Garay JP, et al. Knockin of mutant PIK3CA activates multiple oncogenic pathways. Proceedings of the National Academy of Sciences. 2009;106(8):2835–40.

36. Nussinov R, Tsai CJ, Jang H. A New View of Activating Mutations in Cancer. Cancer Research. 2022 Nov 15;82(22):4114–23.

37. Burke JE, Perisic O, Masson GR, Vadas O, Williams RL. Oncogenic mutations mimic and enhance dynamic events in the natural activation of phosphoinositide 3-kinase p110 (PIK3CA). Proceedings of the National Academy of Sciences. 2012;109(38):15259–64.

38. Jenkins ML, Ranga-Prasad H, Parson MAH, Harris NJ, Rathinaswamy MK, Burke JE. Oncogenic mutations of PIK3CA lead to increased membrane recruitment driven by reorientation of the ABD, p85 and C-terminus. Nat Commun. 2023 Jan 12;14(1):181.

39. Gordus A, Krall JA, Beyer EM, Kaushansky A, Wolf-Yadlin A, Sevecka M, et al. Linear combinations of docking affinities explain quantitative differences in RTK signaling. Molecular Systems Biology. 2009 Jan;5(1):235.

40. Vadas O, Burke JE, Zhang X, Berndt A, Williams RL, Vanhaesebroeck B, et al. Structural basis for activation and inhibition of class I phosphoinositide 3-kinases. Science signaling. 2011;4(195):re2.

41. Gupta S, Ramjaun AR, Haiko P, Wang Y, Warne PH, Nicke B, et al. Binding of Ras to Phosphoinositide 3-Kinase p110α Is Required for Ras-Driven Tumorigenesis in Mice. Cell. 2007;129(5):957–68.

42. Wennström S, Downward J. Role of phosphoinositide 3-kinase in activation of ras and mitogen-activated protein kinase by epidermal growth factor. Molecular and cellular biology. 1999 Jun;19(6):4279–88.

43. Rodriguez-Viciana P, Warne PH, Dhand R, Vanhaesebroeck B, Gout I, Fry MJ, et al. Phosphatidylinositol-3-OH kinase as a direct target of Ras. Nature. 1994 Aug 18;370(6490):527–32.

44. Tsolakos N, Durrant TN, Chessa T, Suire SM, Oxley D, Kulkarni S, et al. Quantitation of class IA PI3Ks in mice reveals p110-free-p85s and isoform-selective subunit associations and recruitment to receptors. Proceedings of the National Academy of Sciences. 2018 Nov 27;115(48):12176–81.

45. Luo J, Field SJ, Lee JY, Engelman J a., Cantley LC. The p85 regulatory subunit of phosphoinositide 3-kinase down-regulates IRS-1 signaling via the formation of a sequestration complex. Journal of Cell Biology. 2005;170(3):455–64.

46. Greshock J, Cheng J, Rusnak D, Martin AM, Wooster R, Gilmer T, et al. Genome-wide DNA copy number predictors of lapatinib sensitivity in tumor-derived cell lines. Molecular Cancer Therapeutics. 2008 Apr 15;7(4):935–43.

47. Burke JE, Williams RL. Synergy in activating class I PI3Ks. Trends in Biochemical Sciences. 2015;40(2):88–100.

48. Yuan TL, Wulf G, Burga L, Cantley LC. Cell-to-cell variability in PI3K protein level regulates PI3K-AKT pathway activity in cell populations. Current Biology. 2011;21(3):173–83.

49. Castel P, Toska E, Engelman JA, Scaltriti M. The present and future of PI3K inhibitors for cancer therapy. Nature Cancer. 2021 Jun 17;180(5):428.

50. Vanhaesebroeck B, Burke JE, Madsen RR. Precision Targeting of Mutant PI3Kα in Cancer by Selective Degradation. Cancer Discovery. 2022 Jan 12;12(1):20–2.

51. Feinberg AP, Levchenko A. Epigenetics as a mediator of plasticity in cancer. Science. 2023;379(6632).

52. Koren S, Reavie L, Couto JP, De Silva D, Stadler MB, Roloff T, et al. PIK3CA(H1047R) induces multipotency and multi-lineage mammary tumours. Nature. 2015 Sep 3;525(7567):114–8.

53. Van Keymeulen A, Lee MY, Ousset M, Brohée S, Rorive S, Giraddi RR, et al. Reactivation of multipotency by oncogenic PIK3CA induces breast tumour heterogeneity. Nature. 2015 Sep 3;525(7567):119–23.

54. Hanker AB, Pfefferle AD, Balko JM, Kuba MG, Young CD, Sanchez V, et al. Mutant PIK3CA accelerates HER2-driven transgenic mammary tumors and induces resistance to combinations of anti-HER2 therapies. Proceedings of the National Academy of Sciences. 2013;110(35):14372–7.

55. Suderman R, Bachman JA, Smith A, Sorger PK, Deeds EJ. Fundamental trade-offs between information flow in single cells and cellular populations. Proceedings of the National Academy of Sciences. 2017;114(22):5755–60.

56. Bivona TG. Dampening oncogenic RAS signaling. Science. 2019 Mar 22;363(6433):1280–1.

57. https://www.sec.gov/Archives/edgar/data/1743881/000119312522146467/d275369dex991.htm

58. Madsen RR, Semple RK. Luminescent peptide tagging enables efficient screening for CRISPR-mediated knock-in in human induced pluripotent stem cells. Wellcome open research. 2019 Jul 11;4:37.

59. Hsiau T, Conant D, Maures T, Waite K, Yang J, Kelso R, et al. Inference of CRISPR Edits from Sanger Trace Data. bioRxiv. 2019;251082.

60. Clement K, Rees H, Canver MC, Gehrke JM, Farouni R, Hsu JY, et al. CRISPResso2 provides accurate and rapid genome editing sequence analysis. Nat Biotechnol. 2019 Mar;37(3):224–6.

61. Kim S, Scheffler K, Halpern AL, Bekritsky MA, Noh E, Källberg M, et al. Strelka2: fast and accurate calling of germline and somatic variants. Nature Methods. 2018;15(8):591– 4.

62. Cingolani P, Platts A, Wang LL, Coon M, Nguyen T, Wang L, et al. A program for annotating and predicting the effects of single nucleotide polymorphisms, SnpEff. Fly (Austin). 2012 Apr 1;6(2):80–92.

63. McLaren W, Gil L, Hunt SE, Riat HS, Ritchie GRS, Thormann A, et al. The Ensembl Variant Effect Predictor. Genome Biology. 2016 Dec 6;17(1):122.

64. Gu Z, Eils R, Schlesner M. Complex heatmaps reveal patterns and correlations in multidimensional genomic data. Bioinformatics. 2016;32(18):2847–9.

65. Ewels PA, Peltzer A, Fillinger S, Patel H, Alneberg J, Wilm A, et al. The nf-core framework for community-curated bioinformatics pipelines. Nature Biotechnology. 2020 Mar 13;38(3):276–8.

66. Dobin A, Davis CA, Schlesinger F, Drenkow J, Zaleski C, Jha S, et al. STAR: Ultrafast universal RNA-seq aligner. Bioinformatics (Oxford, England). 2013;29(1):15–21.

67. Liao Y, Smyth GK, Shi W. FeatureCounts: An efficient general purpose program for assigning sequence reads to genomic features. Bioinformatics. 2014;30(7):923–30.

68. Ritchie ME, Phipson B, Wu D, Hu Y, Law CW, Shi W, et al. limma powers differential expression analyses for RNA-sequencing and microarray studies. Nucleic Acids Research. 2015 Apr 20;43(7):e47–e47.

69. Smid M, Coebergh van den Braak RRJ, van de Werken HJG, van Riet J, van Galen A, de Weerd V, et al. Gene length corrected trimmed mean of M-values (GeTMM) processing of RNA-seq data performs similarly in intersample analyses while improving intrasample comparisons. BMC Bioinformatics. 2018 Jun 22;19(1):236.

70. Benjamini Y, Hochberg Y. Controlling the False Discovery Rate - a Practical and Powerful Approach to Multiple Testing. Journal of the Royal Statistical Society Series B-Methodological. 1995;57:289–300.

71. Norris DM, Yang P, Krycer JR, Fazakerley DJ, James DE, Burchfield JG. An improved Akt reporter reveals intra- and inter-cellular heterogeneity and oscillations in signal transduction. Journal of Cell Science. 2017;130(16):2757–66.

72. Schindelin J, Arganda-Carreras I, Frise E, Kaynig V, Longair M, Pietzsch T, et al. Fiji: an open-source platform for biological-image analysis. Nat Methods. 2012 Jul;9(7):676–82.

73. Dobrzynski M, Jacques MA, Pertz O. Mining single-cell time-series datasets with time course inspector. Bioinformatics. 2019;36(6):1968–9.

74. Schmidt U, Weigert M, Broaddus C, Myers G. Cell Detection with Star-Convex Polygons. In: Frangi AF, Schnabel JA, Davatzikos C, Alberola-López C, Fichtinger G, editors. Medical Image Computing and Computer Assisted Intervention – MICCAI 2018. Cham: Springer International Publishing; 2018. p. 265–73. (Lecture Notes in Computer Science).

75. Tinevez JY, Perry N, Schindelin J, Hoopes GM, Reynolds GD, Laplantine E, et al. TrackMate: An open and extensible platform for single-particle tracking. Methods. 2017 Feb 15;115:80–90.

76. Finck R, Simonds EF, Jager A, Krishnaswamy S, Sachs K, Fantl W, et al. Normalization of mass cytometry data with bead standards. Cytometry Part A. 2013;83A(5):483–94.

77. Zunder ER, Finck R, Behbehani GK, Amir E ad D, Krishnaswamy S, Gonzalez VD, et al. Palladium-based mass tag cell barcoding with a doublet-filtering scheme and single-cell deconvolution algorithm. Nat Protoc. 2015 Feb;10(2):316–33.

78. Hahne F, LeMeur N, Brinkman RR, Ellis B, Haaland P, Sarkar D, et al. flowCore: a Bioconductor package for high throughput flow cytometry. BMC Bioinformatics. 2009 Apr 9;10(1):106.

79. Crowell HL, Zanotelli VRT, Chevrier S, Robinson MD, Bodenmiller B. CATALYST: Cytometry dATa anALYSis Tools [Internet]. Bioconductor version: Release (3.18); 2023 [cited 2023 Dec 19]. Available from: https://bioconductor.org/packages/CATALYST/

80. Stringer C, Wang T, Michaelos M, Pachitariu M. Cellpose: a generalist algorithm for cellular segmentation. Nature Methods. 2020 Dec 14;18(1):100–6.

81. Voliotis M, Perrett RM, McWilliams C, McArdle CA, Bowsher CG. Information transfer by leaky, heterogeneous, protein kinase signaling systems. Proceedings of the National Academy of Sciences. 2014 Jan 21;111(3):E326–33.

82. Foukas LC, Berenjeno IM, Gray A, Khwaja A, Vanhaesebroeck B. Activity of any class IA PI3K isoform can sustain cell proliferation and survival. Proceedings of the National Academy of Sciences of the United States of America. 2010;107(25):11381–6.

83. Qin X, Sufi J, Vlckova P, Kyriakidou P, Acton SE, Li VSW, et al. Cell-type-specific signaling networks in heterocellular organoids. Nature Methods. 2020;17(3):335–42.

84. García-Martínez JM, Alessi DR. mTOR complex 2 (mTORC2) controls hydrophobic motif phosphorylation and activation of serum- and glucocorticoid-induced protein kinase 1 (SGK1). The Biochemical journal. 2008;416(3):375–85.

